# Human cerebral organoids model tumor infiltration and migration supported by astrocytes in an autologous setting

**DOI:** 10.1101/2025.01.29.635456

**Authors:** Esther Schickel, Tamara Bender, Leon Kaysan, Simone Hufgard, Margot Mayer, David R. Grosshans, Christiane Thielemann, Insa S. Schroeder

## Abstract

Efforts to achieve precise and efficient tumor targeting of highly malignant brain tumors are constrained by the dearth of appropriate models to study the effects and potential side effects of radiation, chemotherapy, and immunotherapy on the most complex human organ, the brain. We established a cerebral organoid model of brain tumorigenesis in an autologous setting by overexpressing c-MYC as one of the most common oncogenes in brain tumors. GFP^+^/c-MYC^high^ cells were isolated from tumor organoids and used in two different culture approaches: assembloids comprising of a normal cerebral organoid with a GFP^+^/c-MYC^high^ tumor sphere and co-culture of cerebral organoid slices at air-liquid interface with GFP^+^/c-MYC^high^ cells. GFP^+^/c-MYC^high^ cells used in both approaches exhibited tumor-like properties, including overexpression of the c-MYC oncogene, high proliferative and invasive potential, and an immature phenotype as evidenced by increased expression of Ki-67, VIM, and CD133. Organoids and organoid slices served as suitable scaffolds for infiltrating tumor-like cells. Using our highly reproducible and powerful model system that allows long-term culture, we demonstrated that the migratory and infiltrative potential of tumor-like cells is shaped by the environment in which glia cells provide support to tumor-like cells.

## Introduction

Malignant brain tumors present a significant societal burden, as no effective therapeutic options are currently available despite extensive research efforts in surgery, chemotherapy, immunotherapy, and radiotherapy. For instance, the prognosis for glioblastoma patients remains poor, with a median survival of approximately 15 months [1, 2]. Pediatric patients are particularly affected, as they do not respond well to most of the standard treatments [3–5]. The reconsideration of treatment modalities necessitates the development of more appropriate human tissue and organ models than those previously described [6]. For instance, a monolayer of tumor cells in a dish or trans-well insert can be employed for studies of tumor migration [7, 8], but not infiltration due to the lack of a three-dimensional environment. Furthermore, the loss of cellular diversity resulting from clonal selection during cell propagation and increased genetic variation within cell lines represent additional drawbacks of such cultures [9]. This, in turn, diminishes their physiological relevance. Animal models, such as rats and mice, can be used to inject tumor cells, to be xenografted with a patient’s tumor tissue, or allografted with genetically modified cells [10, 11]. However, these models fail to accurately recapitulate the situation in humans due to species differences in the genetic background, morphology, anatomy, and metabolic states [12]. Regarding the potential use of human cells, one or more cell types can be aggregated and maintained as uniform spheres. However, they lack specific organization and complexity, as observed in organoids. Conversely, cerebral organoids comprising diverse cell types, including neuronal and glial cells derived from pluripotent cells, represent a promising platform for the generation of human brain tumors. A variety of approaches have been employed to achieve this goal, including the use of patient-derived explants of brain tumors cultured in scaffolds or matrices or dissociated tumor cells that can self-arrange into patient-derived organoids [13, 14]. However, such models are unfortunately not suitable for long-term culture due to the loss of tumor heterogeneity during their propagation [15]. Further approaches described in the literature include the co-culture of cerebral organoids with primary tumor cells or cell lines [16]. However, the genetic and epigenetic landscapes of each entity differ, which presents a challenge for the formation of the tumor in its original surrounding. Genetic manipulations of oncogenes and tumor suppressor genes can be induced in the organoid itself by tools such as CRISPR/Cas9 and/or transposase to enable the formation of the tumor in its original surrounding [17, 18]. Here we present an adaptation and combination of existing protocols, resulting in an autologous brain tumor model where long-term culture is possible. For this purpose, normal cerebral organoids were generated using human pluripotent stem cells and a protocol for their 3D organization and differentiation into neuronal and glial cells [19, 20]. In addition, tumor organoids containing genetically modified cells were generated using the SB transposon system for the induction of c-MYC oncogene overexpression [18]. During the culture period, organoids undergo growth and typically develop a necrotic core due to inadequate supply of oxygen and nutrients from the medium [21]. To circumvent this issue, we prepared 300 µm organoid slices and cultured them at an air-liquid interface, which provided a dual supply of nutrients from the medium and oxygen from the air. This enabled us to achieve long-term culture [22] beyond 100 days without the occurrence of necrosis. To generate a reproducible tumor model in an autologous setting, we either co-cultured organoid slices with a defined number of tumor-like cells isolated from sister organoids using fluorescence-activated cell sorting (FACS) based on the presence of GFP signal or used them to generate tumor-spheres that were able to fuse with whole sister organoids to form assembloids. Tumor-like cells interacted with normal cells from the organoids or organoid slices, e.g. astrocytes, that provided a support for their proliferation, migration/infiltration.

## Results

We established and characterized cerebral organoids and organoid slices as brain tumor models by combining previously published protocols [19, 20] as a basis for the introduction of genetic modifications to generate tumor-like cells [18] and for the culture at the air-liquid interface (ALI) [22] to allow long-term maintenance of the system. This is shown schematically in the workflow (Figure 1). The characteristics of brain-like organoids and organoid slices were evaluated, including cell composition, cell interactions, and dynamics that enable processes and functions in these structures. Tumor-like properties of transfected, c-MYC^+^/GFP^+^ cells were confirmed, including oncogene overexpression, immature or stem cell-like properties, highly proliferative and invasive potential of genetically modified cells used for assembloids with brain organoids or co-culture with organoid slices.

**Figure 1.**
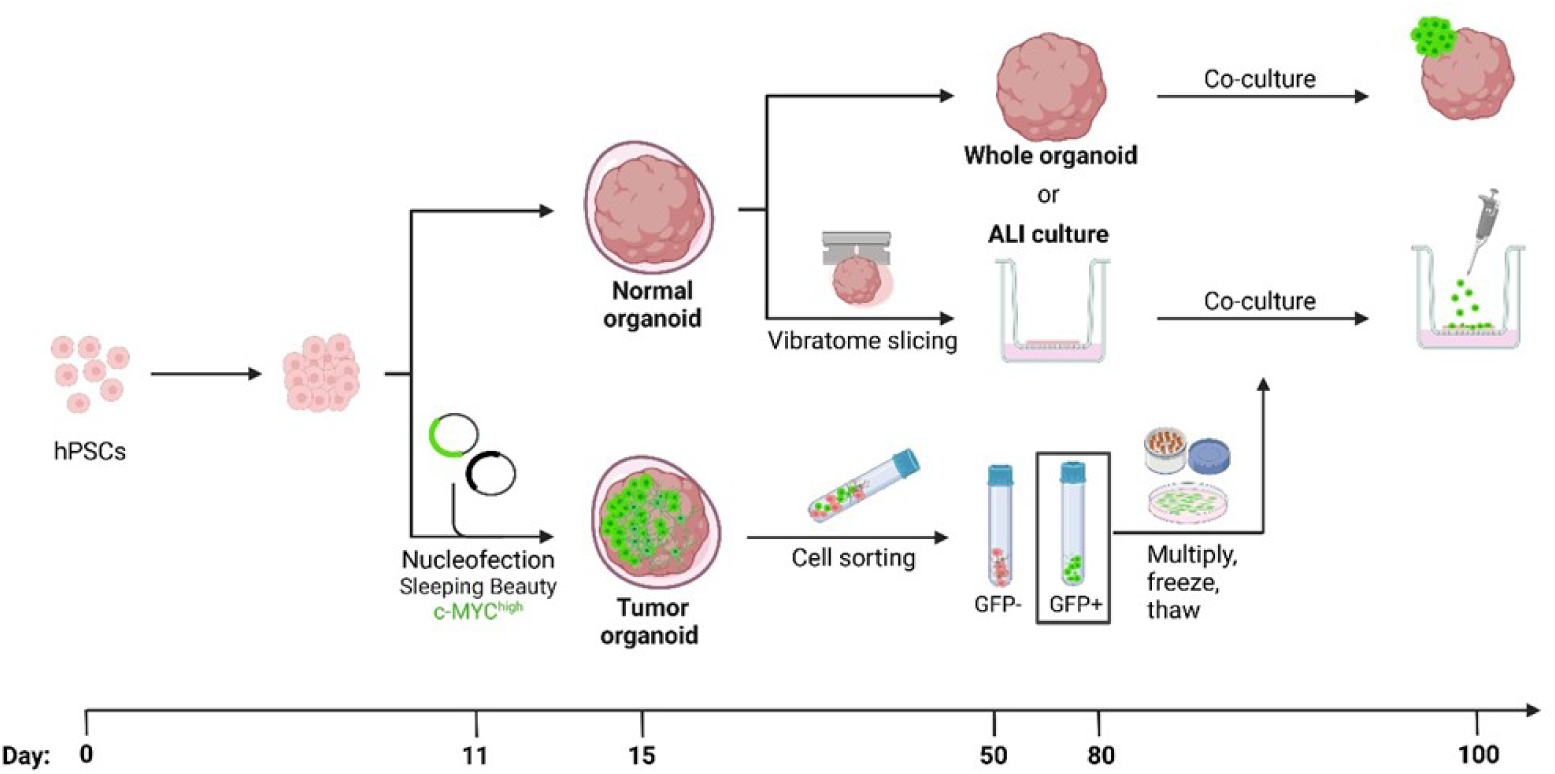
Workflow for generation of the tumor model: tumor spheres assembled with cerebral organoid and cerebral organoid slices co-cultured with tumor-like cells at air liquid interface. Generation of embryoid bodies using human pluripotent stem cells (hPSCs). Nucleofection of organoids on day 11 of culture using Sleeping Beauty Transposon System for overexpression of c-MYC oncogene with co-expressed GFP. Embedding of organoids in Matrigel on day 15 of culture. Normal organoids cultured as whole entities or organoid slices prepared using vibratome at ∼ day 50 and maintained on cell culture inserts. Normal organoids or organoid slices cultured in parallel with nucleofected organoids used for isolation of genetically modified cells based on the presence of GFP signal using fluorescence-activated cell sorting at ∼day 50 for co-culture wit organoid slices or at ∼and 80 for generation of tumor spheres to be assembled with whole organoids. Positively selected (GFP^+^) cells can be multiplied, frozen and thawed before used for either generation of spheres to be assembled with whole organoid or as single

### Cerebral organoids exhibit dynamic landscape of proliferation and maturation

Cerebral organoids generated from hESCs showed an increase in size during the culture period as evidenced by the measured organoid circular area. This was attributed to intense cell proliferation as evidenced by the increase in the measured organoid circular area and from Ki-67 positive staining up to approximately day 50 in culture, after which no further increase in organoid size and little to no Ki-67 signal was observed (Figure 2A and 2B). Notably, different cell types were present in the organoids at the same time, but the protein expression of their specific markers and their spatial distribution was dependent on the organoid’;s age (Figure 2B shows spatial distribution while Figure S1 displays quantification of the IF signal). For instance, early progenitors expressing nestin were present at approximately day 25 of culture with distribution throughout the organoid. Beyond day 100, nestin-expressing cells were predominantly found at the outer edge of the organoid. Neuronal markers such as MAP2 and DCX were also detectable before day 50 of differentiation and were expressed throughout the organoid. Their expression peaked around day 50, after which it slightly decreased and the MAP2 and DCX positive cells were predominantly found at the outer edge of the organoid, similar to the spatial distribution of nestin. The marker of axons (SMI312) showed an increase in expression around day 50 of culture, which continued beyond day 100 with no further significant increase in expression intensity. An increase in the abundance of astrocytes, as indicated by the expression of GFAP, started around day 100 and continued until day 150 at the outer edge of the organoid. The marker for cell junctions (CX43) showed a similar expression pattern, however, here the increase in expression started slightly earlier at about day 75 but also reached its plateau at d150 (Figure 2B and Figure S1). In addition, the pre-synaptic markers SYN1 and VAMP2 were localized near neuronal cells (MAP2^+^ or SMI312^+^) at the outer edge of organoids where astrocytes (GFAP^+^) were also found at ∼d100-125 (Figure 2C-D). The presynaptic marker VAMP2, primarily localized to the synaptic vesicles in the axonal terminal and involved in neurotransmitter release, was observed in close proximity to the postsynaptic marker HOMER (Figure 2E, Figure S2).

**Figure 2.**
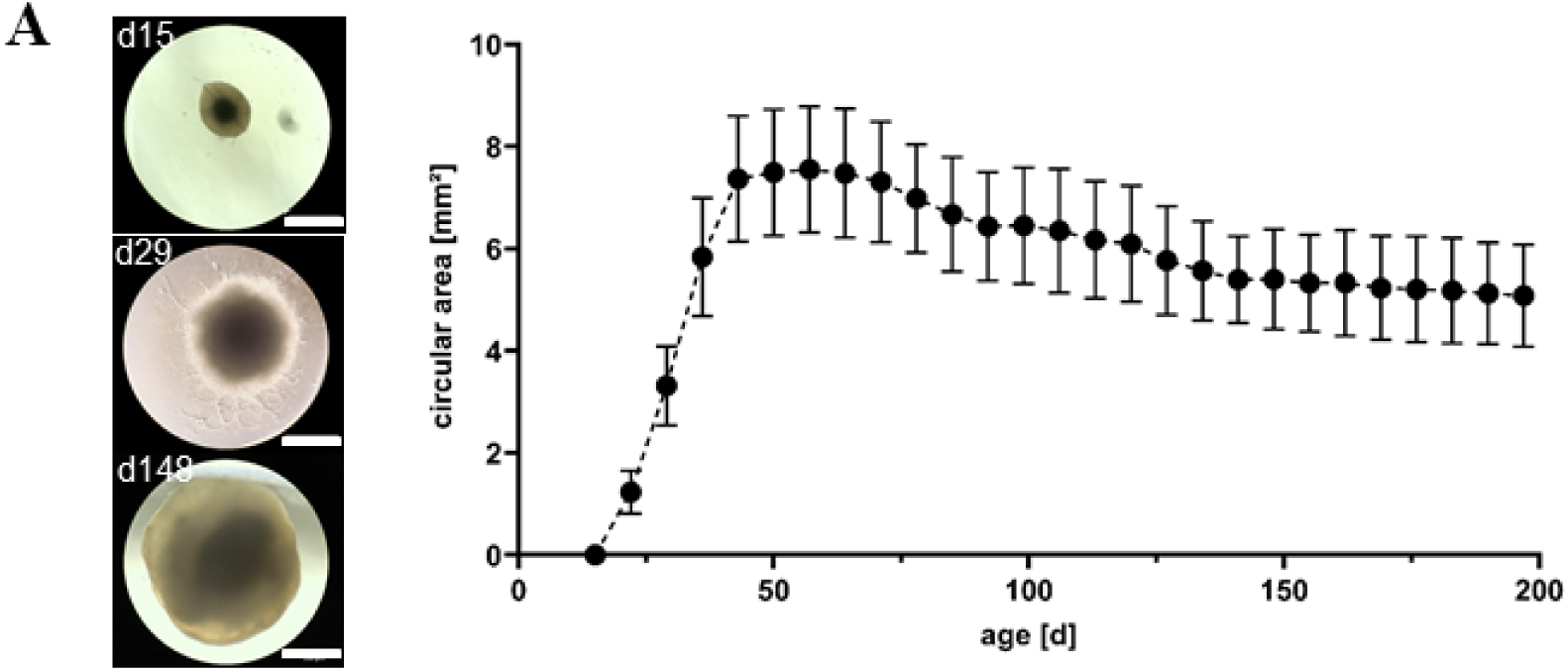
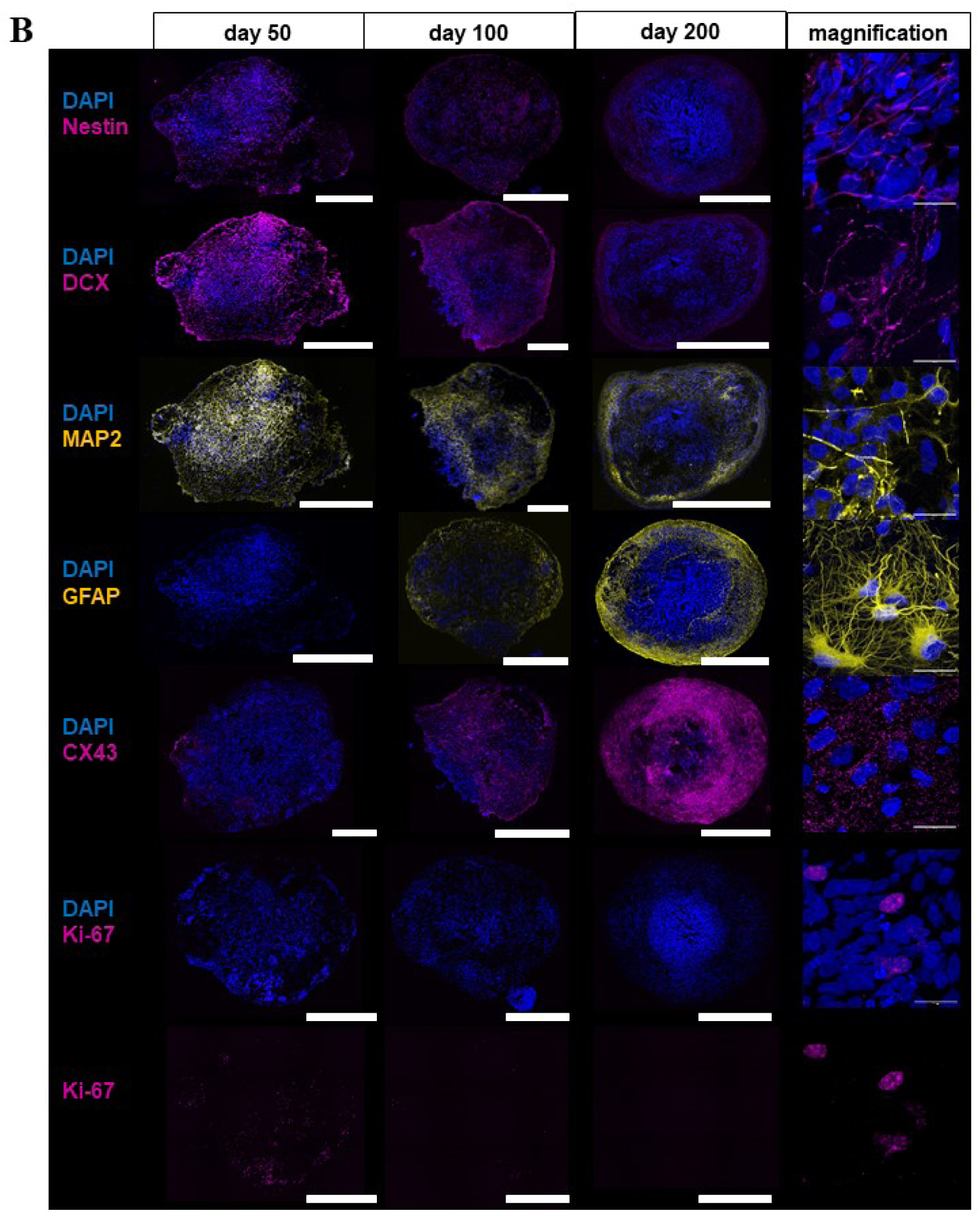
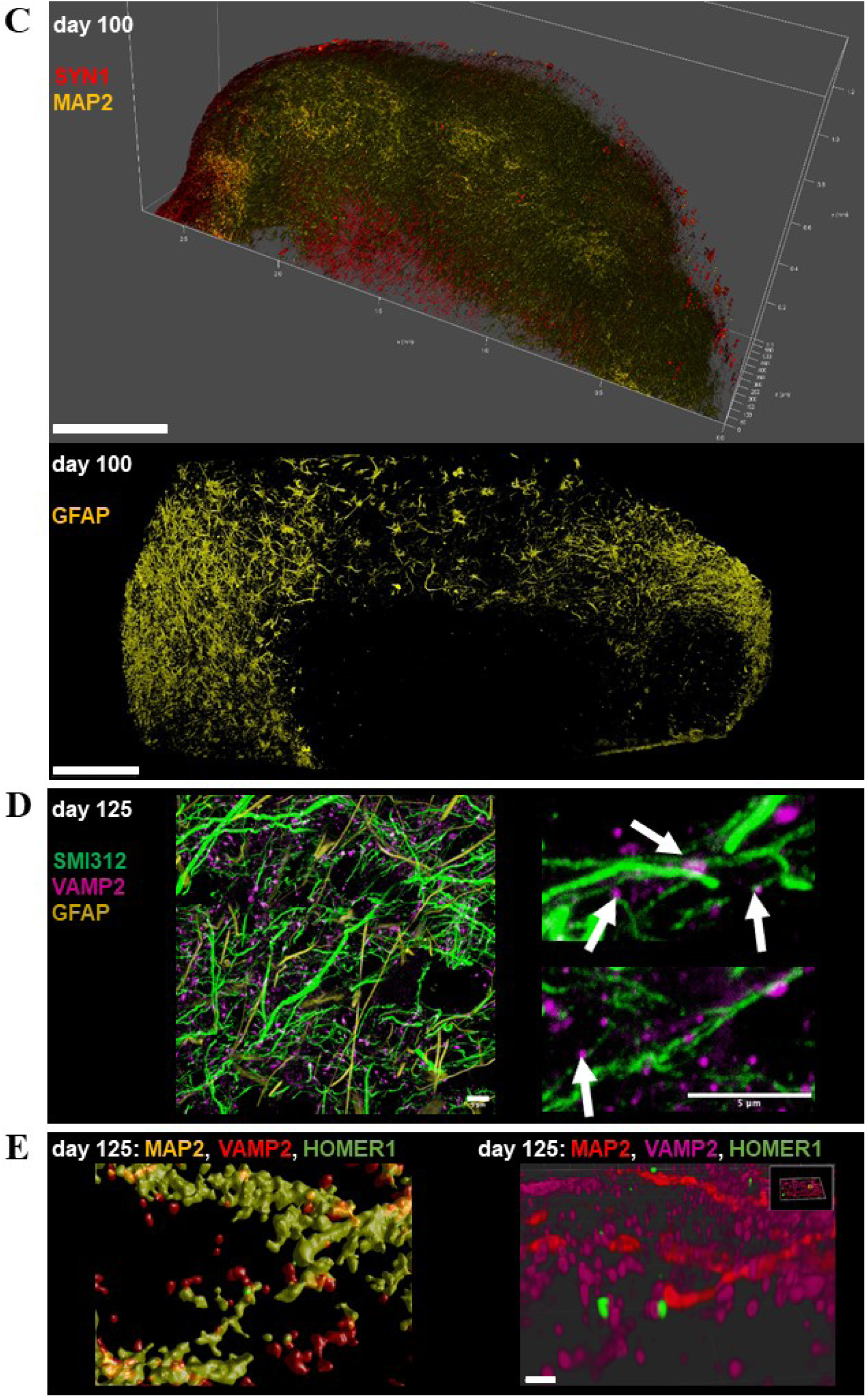
Proliferation and maturation in cerebral organoids. **A)** Representative microscopy images of organoids at d15, d29 and d148 of the culture (left). The organoid size measured as circular area [mm^2^] for organoids between d15 and d200 104 organoids per experiment (n = 24-104). Scale bar: 1000 µm. **B)** Representative immunofluorescence staining of nestin (magenta), DCX (magenta), MAP2 (yellow), GFAP (yellow), CX43 (magenta), and Ki-67 (magenta) for organoids at d50, d100 and d200 of the culture, respectively. Scale bar: 1000 µm. Magnification for nestin at d50, DCX at day 100, MAP2 at d100, GFAP at d150, CX43 at d200 and Ki-67 at d50. Scale bar: 20 µm. **C)** Representative immunofluorescence staining of SYN1 (red) in combination with MAP2 (yellow) (700 µm z-stack), and of GFAP (yellow) (overview images) at d100. Scale bar: 500 µm for SYN1 and MAP2, 200 µm for GFAP staining. **D)** Magnification image of representative immunofluorescence staining of SMI312 (green), VAMP2 (magenta; indicated with white arrows (right image) and GFAP (yellow) at d125. Scale bar: 5 µm. **E)** Magnification image of representative immunofluorescence staining of HOMER1 (green), VAMP2 (red), MAP2 (yellow) at d125 after deconvolution (right), and surface rendering (left). Scale bar: 3 µm.

### Cerebral organoids and organoid slices are robust models that exhibit brain-like features

An increase in organoid size during the culture period was correlated with the occurrence of cell death in the organoid interior, indicated by shrinked and fragmentated nuclei in DAPI staining and measured as lactate dehydrogenase (LDH) release (Figure 3A). This was due to an inadequate supply of oxygen and nutrients, which is a limiting factor for prolonged culture. In contrast to whole organoids, organoids cut into 300 µm thick slices showed mostly intact nuclei and significantly less extracellular LDH than whole organoids of the same age while maintaining the original 3D architecture. The abundance of proliferative cells (Ki-67^+^) was low in whole organoids at day 100 of culture and comparable to that in organoid slices of the same age (Figure S3A-C). The occurrence of apoptosis, as evidenced by caspase-3 active staining at day 100 was very low in both culture systems (Figure S3A-C).

**Figure 3.**
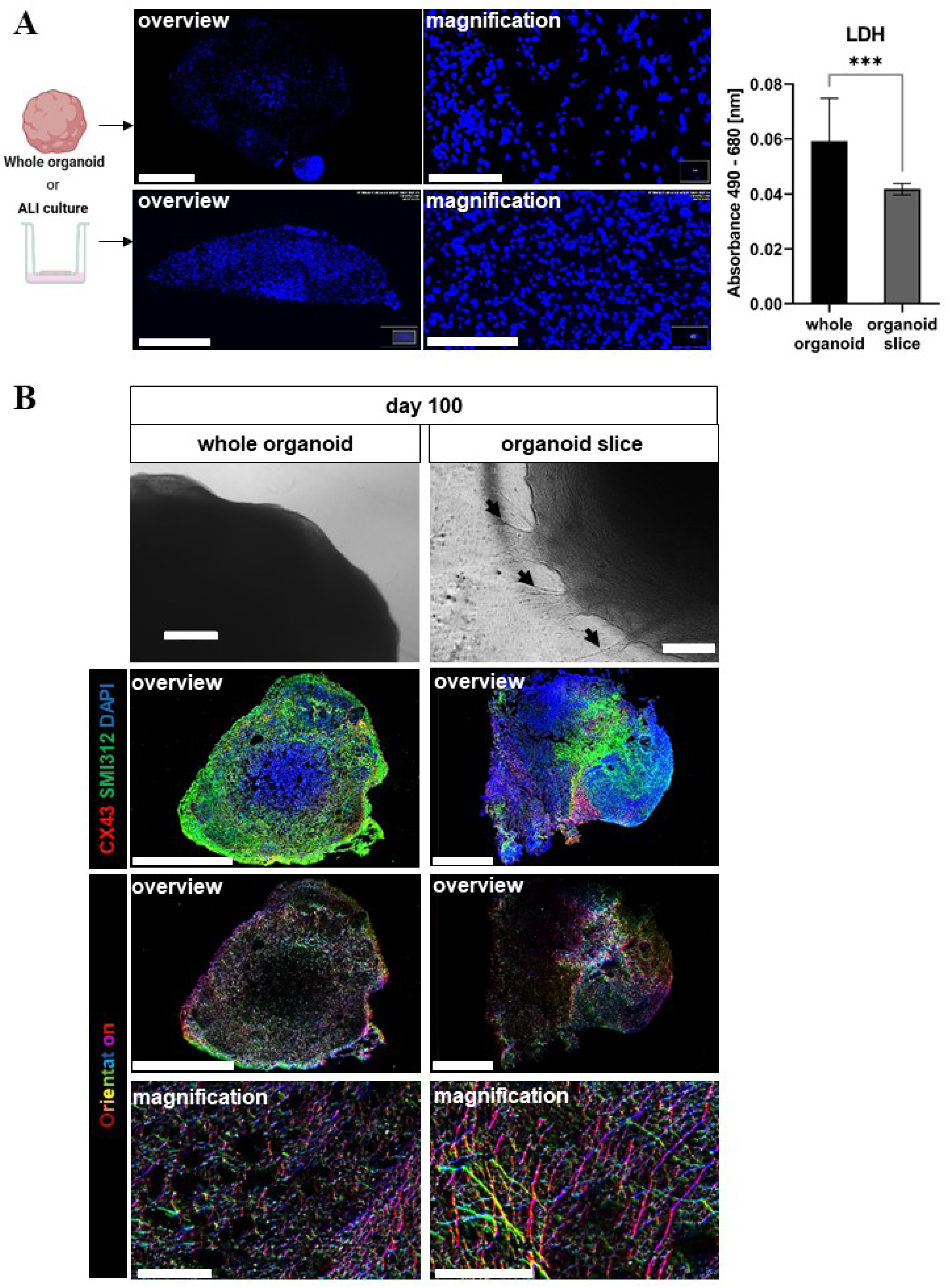
Whole organoids versus organoid slices culture at the air-liquid interface (ALI) at day 100 of the culture. **A)** Representative DAPI staining of whole organoids cultured in medium and in organoid slices cut into 300 µm thick sections cultured on cell culture inserts at the ALI in overview and magnification. Scale bar: 1000 µm in overview and 200 µm in magnification (left). Lactate dehydrogenase (LDH) level measured in whole organoids cultured in medium and in organoid slices cut into 300 µm thick sections cultured on cell culture inserts at the ALI. Data are presented as mean ± SD for three to six p<0.0001. Statistical analysis was done using unpaired t-test (right). **B)** Representative phase contrast image of the whole organoid with smooth edges, and organoid slice showing protrusions extending from the main body of the organoid slice (black arrow). Scale bar: 320 µm. Representative immunofluorescence staining of CX43 (red) in combination with SMI312 (green) and orientation analysis for SMI312 staining (pseudocolor). Scale bar: 1000 µm in overview and 100 µm in magnification.

Organoid slices cultured at the air-liquid interface (ALI) showed axon tracts extending away from the main body of the slice that grew in width and size over time (Figure 3B). Organoid slices were maintained for 100 days and beyond and continued to display brain-like features, including dynamic and organized axonal networks, as evidenced by staining against an axonal marker (SMI312) that revealed specific orientations of axon fibers, in contrast to the lack of directionality in orientation analysis observed in age-matched whole organoids. Furthermore, intercellular networks were confirmed by staining for gap junction protein (CX43) (Figure 3B). At day 100, organoids and organoid slices exhibited a heterogeneous cellular composition (Figure S4), including glial precursor cells (GLAST^+^ and PDGFRα^+^), neurons (MAP2^+^), and radial glia and mature astrocytes (GFAP^+^). While there was no significant difference in the expression of GFAP between the two culture systems at the mRNA level, its spatial distribution in whole organoids predominantly at the outer area was different from organoid slices where it was more equally distributed. Furthermore, organoid slices showed a higher expression of MAP2 and lower expression of vimentin (VIM, an intermediate filament protein expressed by immature astrocytes [23], compared to whole organoids at the mRNA level. Notably, IF and bulk RNA sequencing analyses revealed that expression of PDGFRα, a marker of oligodendrocytes [24], and glia cell markers in general, were more prominent in organoid slices than in whole organoids, suggesting that culture at an air-liquid interface leads to enhanced cell maturation (Figure S4A-B, Figure S5). Such organization and composition enabled processes and functions such as myelination, which is mediated by myelin basic protein (MBP) and spontaneous cell activity. Although the expression of MBP was significantly higher at the mRNA level in organoid slices compared to organoids at day 100, its protein expression was rather low in both model systems, as confirmed by immunofluorescence staining (Figure S4B), suggesting that myelination is still in an early phase. Nevertheless, spontaneous activity of cells in organoid slices was observed in calcium signal recordings (Figure S4C). Finally, culture at ALI allowed us to achieve long-term culture beyond one year (Figure S6).

### Genetically modified cells of brain organoids display tumor-like properties

To generate a reproducible tumor model comprising genetically modified tumor-like cells fused as a tumor sphere with whole organoids (assembloid) or co-cultured with organoid slices, we first optimized the nucleofection procedure (including plasmid concentration, and nucleofection program) (Figure S7). This was of crucial importance to preserve organoid integrity allowing for optimal propagation of tumor-like cells. Nucleofection of organoids with vectors carrying SB Transposase and c-MYC (Figure S7A) oncogene coupled to GFP resulted in the first appearance of GFP^+^ cells within 1 week after nucleofection, and in most cases their relatively rapid overgrowth of organoids as evidenced by the increase in GFP signal during the culture time (Figure 4A), which was also a limiting factor for a prolonged culture of nucleofected organoids. In addition, the randomness of nucleofection resulted in high variability of c-MYC overexpression observed as different GFP signal intensities between organoids (Figure S7E). To enhance the reproducibility of experiments, we isolated genetically modified cells from nucleofected organoids based on the presence of the co-expressed GFP, which was identified by fluorescence-activated cell sorting (FACS) and confirmed by immunofluorescence (IF) staining (Figure 4B-C). The cells could be passaged and/or cryopreserved for later use. As shown in Figure 4C, these GFP^+^/c-MYC^high^ cells exhibited high proliferative capacity (Ki-67^+^) and an immature phenotype (Sox2^+^), with the expression of an axonal marker (SMI312) to some extent and almost no detectable expression of the glial progenitor marker PDFGRα. In addition, they displayed invasive properties (VIM^+^) and a (cancer)-stem-cell-like phenotype (CD133^+^) (Figure 4C). qPCR analyses comparing the mRNA levels in controls, nucleofected organoids and isolated GFP^+^/c-MYC^high^ cells revealed a significant increase in the expression of c-MYC in nucleofected organoids and in isolated GFP^+^ cells (Figure 4D). GFP^+^ cells exhibited a significant increase in the expression of p53, Ki-67, CD133, GLS, and SNAI1 at mRNA level when compared to control organoids. However, there was no change in the expression of these markers between control and nucleofected whole organoids. Expression of the tumor suppressor markers NF1 and PTEN was significantly decreased in nucleofected organoids when compared to control organoids and GFP^+^ cells. Conversely, GFP^+^ cells exhibited no significant difference in the expression of NF1 and PTEN compared to control organoids (Figure 4D). Notably, certain proteins, such as c-MYC and GFP, exhibited considerable heterogeneity in the signal intensity observed in IF staining. GFP^+^ cells exhibited a strong albeit not significant decrease of neuronal and glial markers compared to the control organoids due to this heterogeneity (Figure S8).

**Figure 4.**
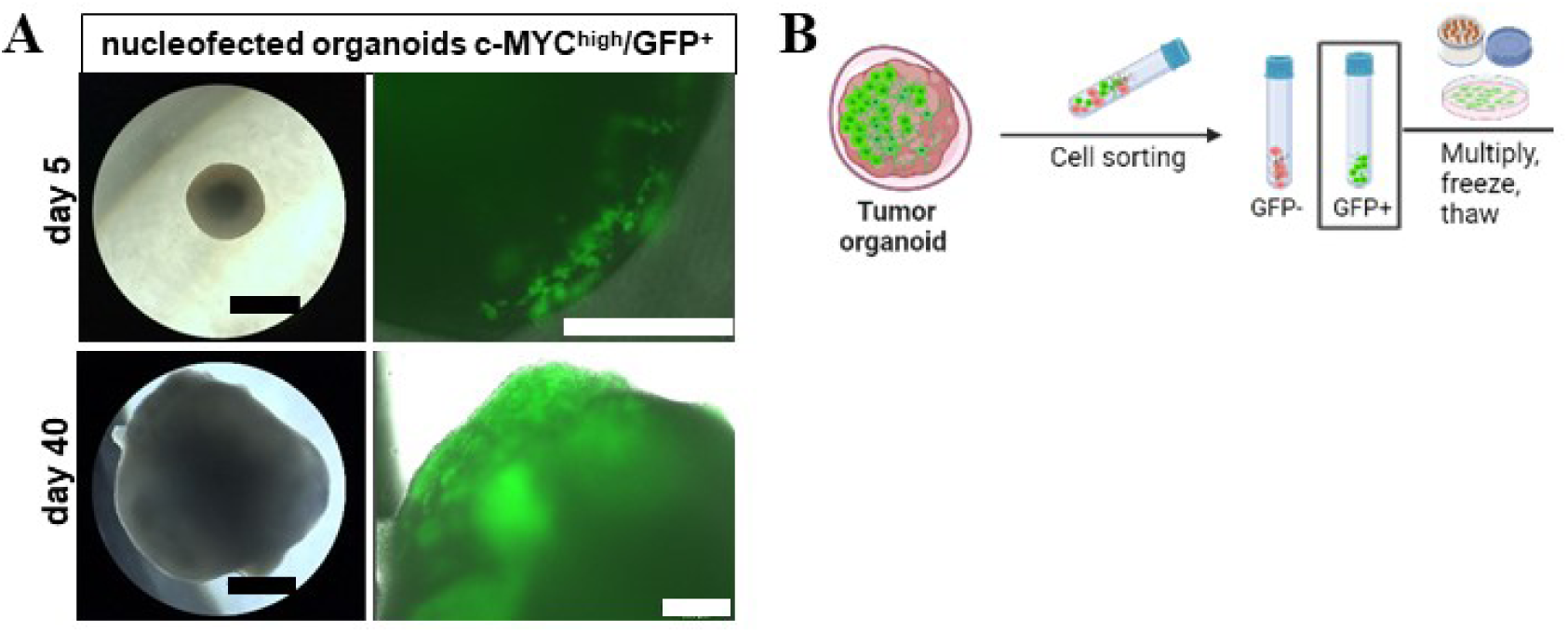
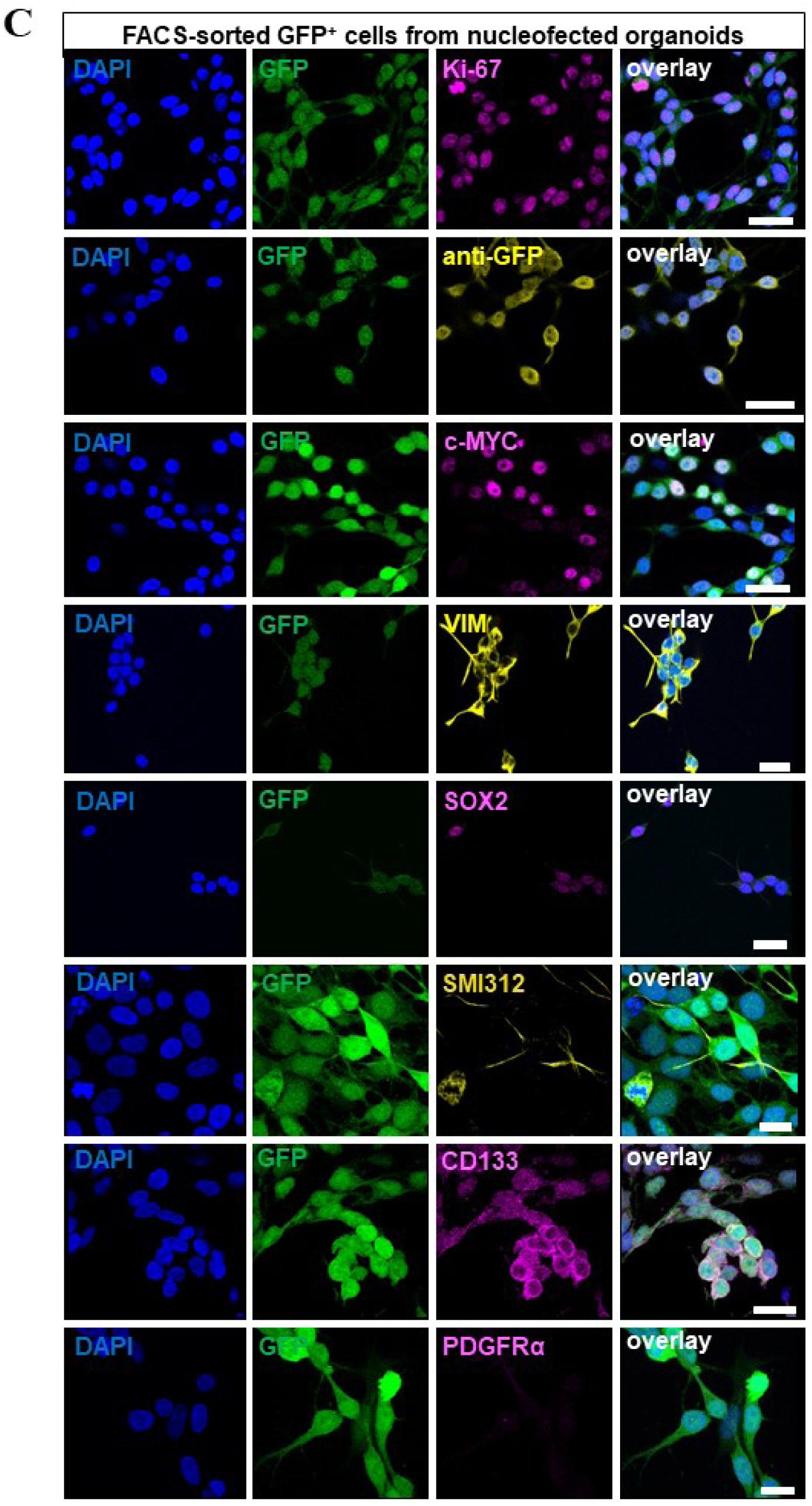
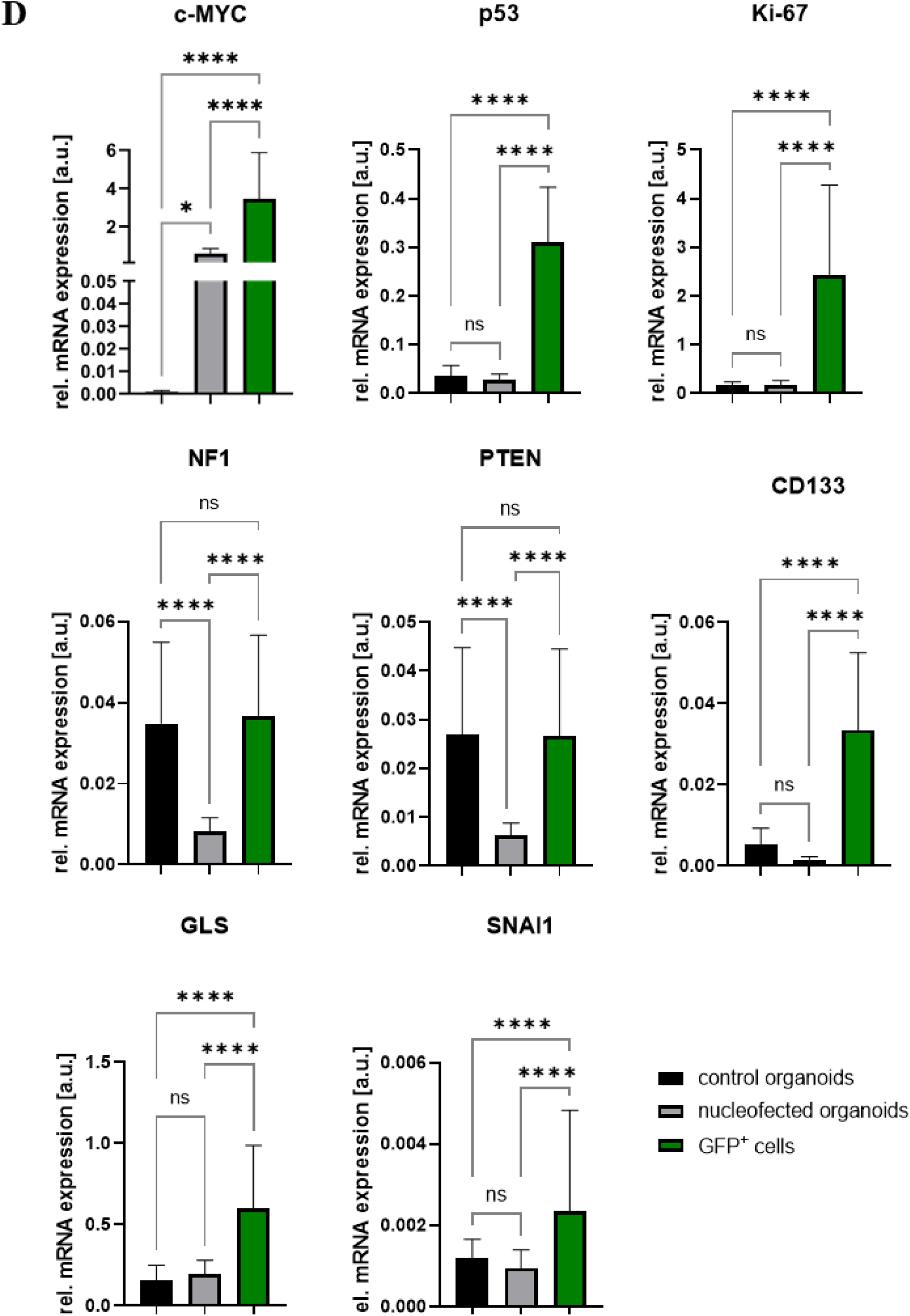
Genetically modified tumor-like cells generated in organoids show tumor-like properties. **A)** Representative microscopic images of organoids at d5 and d40 after nucleofection. Scale bar: 930 µm left and 330 µm right. **B)** GFP^+^/c-MYC^high^ cells isolated from nucleofected organoids based on the presence of GFP that can be further propagated as adherent cells or cryopreserved for later use. **C)** Representative immunofluorescence staining of Ki-67 (magenta), anti-GFP (yellow), SOX2 (magenta), VIM (yellow), CD133 (magenta), c-MYC (magenta), PDGFRα (magenta), and SMI312 (yellow) for GFP^+^ cells (green). Scale bar: 25 µm. **D)** Relative mRNA expression of c-MYC, p53, Ki-67, NF1, PTEN, CD133, GLS and SNAI1 in GFP^+^ cells compared to control (sham-nucleofected) and nucleofected whole organoids. Data are presented as mean ± SD for three independent experiments (N = 3) and three organoids per experiment (n = 3), * p<0.05, ** p<0.01, *** p<0.001, **** p<0.0001. Statistical analysis was done using 2-way mixed ANOVA with Tukeýs post-test.

### Cerebral organoids and organoid slices serve as scaffolds for autologous tumor-like cells

To increase comparability and reproducibility of the tumor model, GFP^+^/c-MYC^high^ cells were aggregated to form spheres and fused with normal sister organoids (assembloids).

The tumor sphere part within the assembloids exhibited an increased cell density from day 9 after fusion compared to the tumor spheres cultured alone (Figure 5A). Through the fusion, the GFP^+^/c-MYC^high^ cells of the tumor sphere encountered contact to astrocytic cells (GFAP^+^) and neurons (MAP2^+^) from the outer edge of the organoid (Figure 5B-C). The protrusions of astrocytic cells (GFAP^+^) were observed to spread from the interaction site into the tumor sphere, while this was observed less extensively for the neurons (MAP2^+^) (Figure 5C). In turn, the GFP^+^ cells from the tumor sphere demonstrated the ability to migrate into the organoid (Figure 5D). The migratory potential of GFP^+^/c-MYC^high^ cells from the sphere was evident by the presence of GFP signal and IF staining in the organoid, particularly at sphere-organoid interface, with a small number of GFP^+^/c-MYC^high^ cells that were observed in deeper areas of the organoid (Figure 5C). Moreover, individual migrating GFP^+^ cells seemed to closely interact with astrocytic cells (GFAP^+^), which surrounded them with their protrusions (Figure 5D). In the co-culture of organoid slices and single tumor-like cells added on top of the slice, we observed an increase in GFP signal over time, resulting in overgrowth of the organoid slice (Figure 6A). GFP^+^/c-MYC^high^ cells were predominantly found near areas of the slice with increased GFAP and GLAST expression, although there was no or only little co-localization of the signals (Figure 6B-C). GFP^+^ cells, despite their proximity to areas with increased expression of neuronal markers (SMI312 and MAP2), were typically spatially segregated from those areas (Figure 6D). In addition, GFP^+^ cells were observed in deeper layers of organoid slices (Figure 6E), confirming their potential for infiltration and invasion. Cells in both tumor models retained proliferative and immature characteristics, as evidenced by c-MYC overexpression and increased Ki-67 expression, and decreased expression of MAP2 and GFAP compared to their respective controls (Figure 7A-D). The expression of p53 was higher in the co-culture than in the control (Figure 7E), while the expression of NF1 and PTEN was decreased in both tumor models compared to normal organoids or slices (Figure 7F-G). This reflects the situation in the whole nucleofected organoids (Figure 7). In contrast to its expression in GFP^+^ cells, CD133 showed a decreased expression in the co-culture compared to the control (Figure 7H). There was no discernible difference in the expression of GLS between the control and co-culture, while the expression of SNAI1, a master regulator of epithelial to mesenchymal transition [25], was increased in the co-culture, but not in the assembloids (Figure 7I-J). Both tumor models showed a significant increase in necrosis-associated LDH level in the cell culture medium compared to their respective controls (Figure 7K).

**Figure 5.**
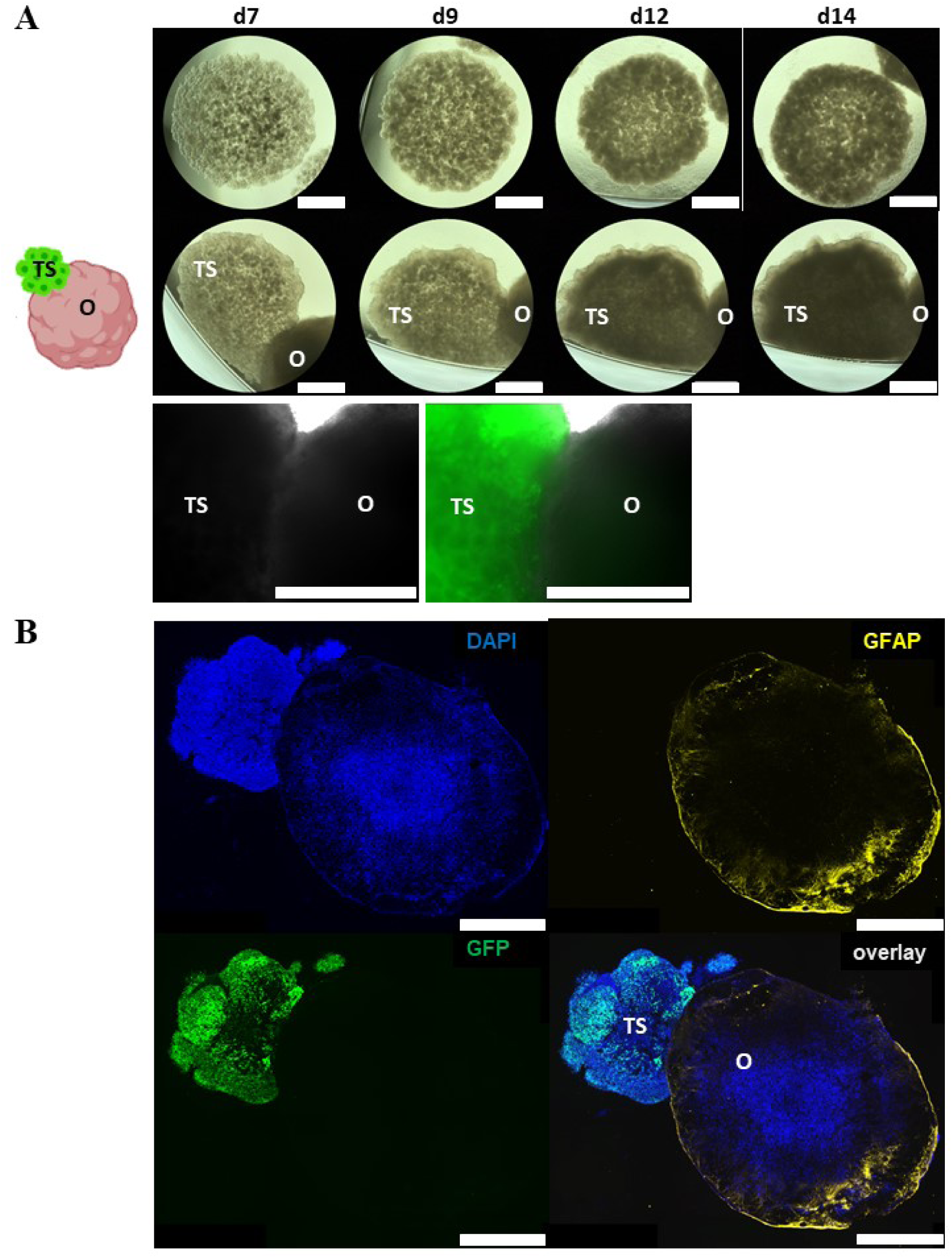
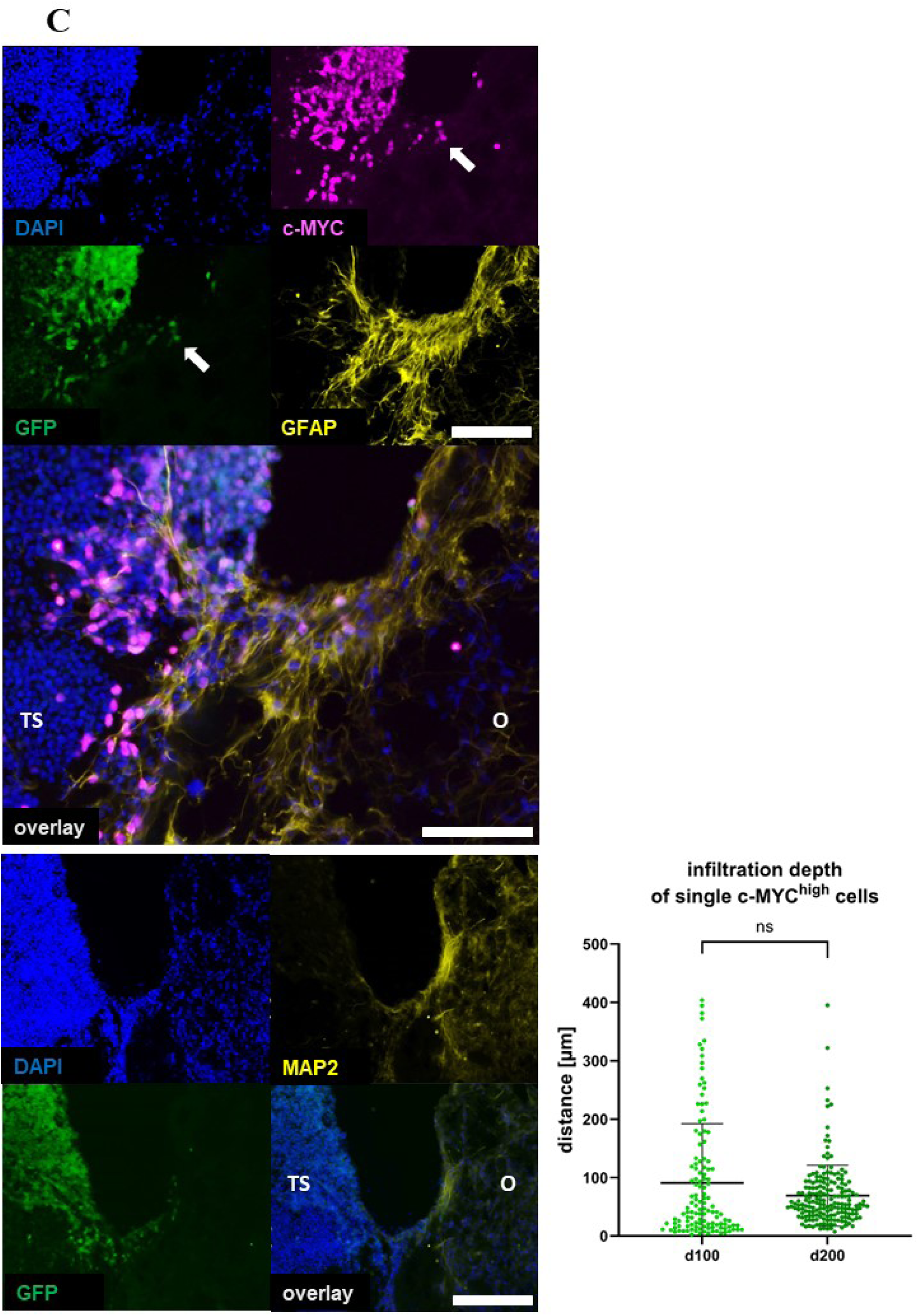
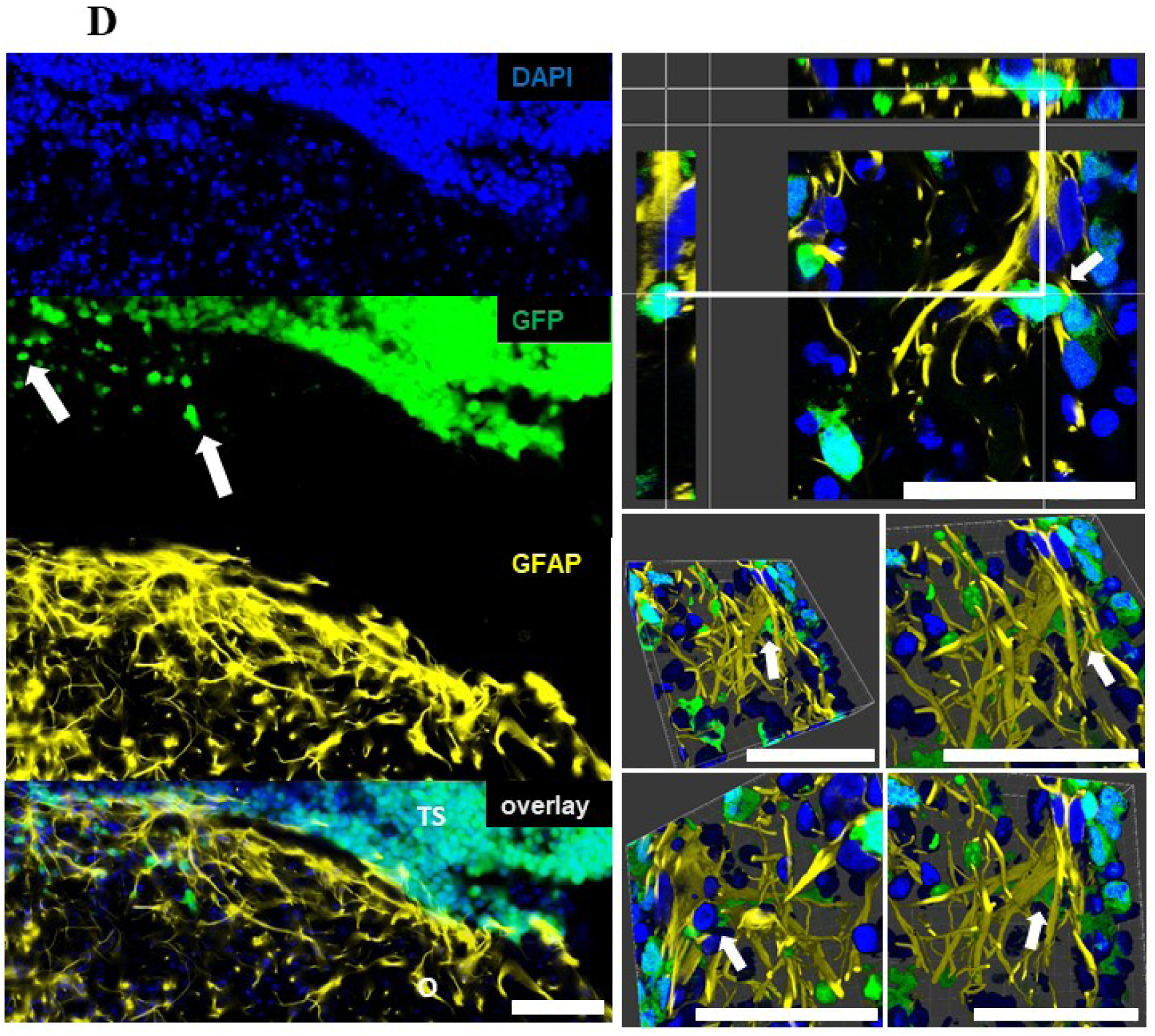
Genetically modified tumor-like cells assembled with whole organoids or co-cultured with organoid slices. **A)** Spheres generated using GFP^+^ tumor-like cells alone and after fusion with the whole organoid. **B)** Representative immunofluorescence staining of GFAP (yellow) for organoid fused with tumor-sphere with GFP^+^ cells (green) in 30 μm thick cryosection. Scale bar: 1000 µm. **C)** Representative immunofluorescence staining of c-MYC (magenta) for GFP^+^ cells (green) in the tumor sphere and of GFAP (yellow) or MAP2 (yellow) in the organoids at their interaction site in 30 μm thick cryosection. Scale bar: 100 µm. Infiltration depth of GFP^+^/c-MYC^high^ cells measured as a distance from the interaction site with organoids at day 100 and day 200 of the culture. Data are presented as mean + SD for three independent experiments (N = 3) and two to three organoids per experiment (n = 2-3), while each point represents the measurement of one single GFP^+^ c-Myc^high^ cell (125 cells at d100 and 185 cells at d200). Statistical analysis was done using Mann Whitney test. **D)** Representative immunofluorescence staining of GFAP (yellow) for the astrocytes of the organoid in the interaction with GFP^+^ cells (green) in the 3D image of 30 μm thick cryosection generated using z-stack. Scale bar: 50 µm (right side) and 100 µm (left side).

**Figure 6.**
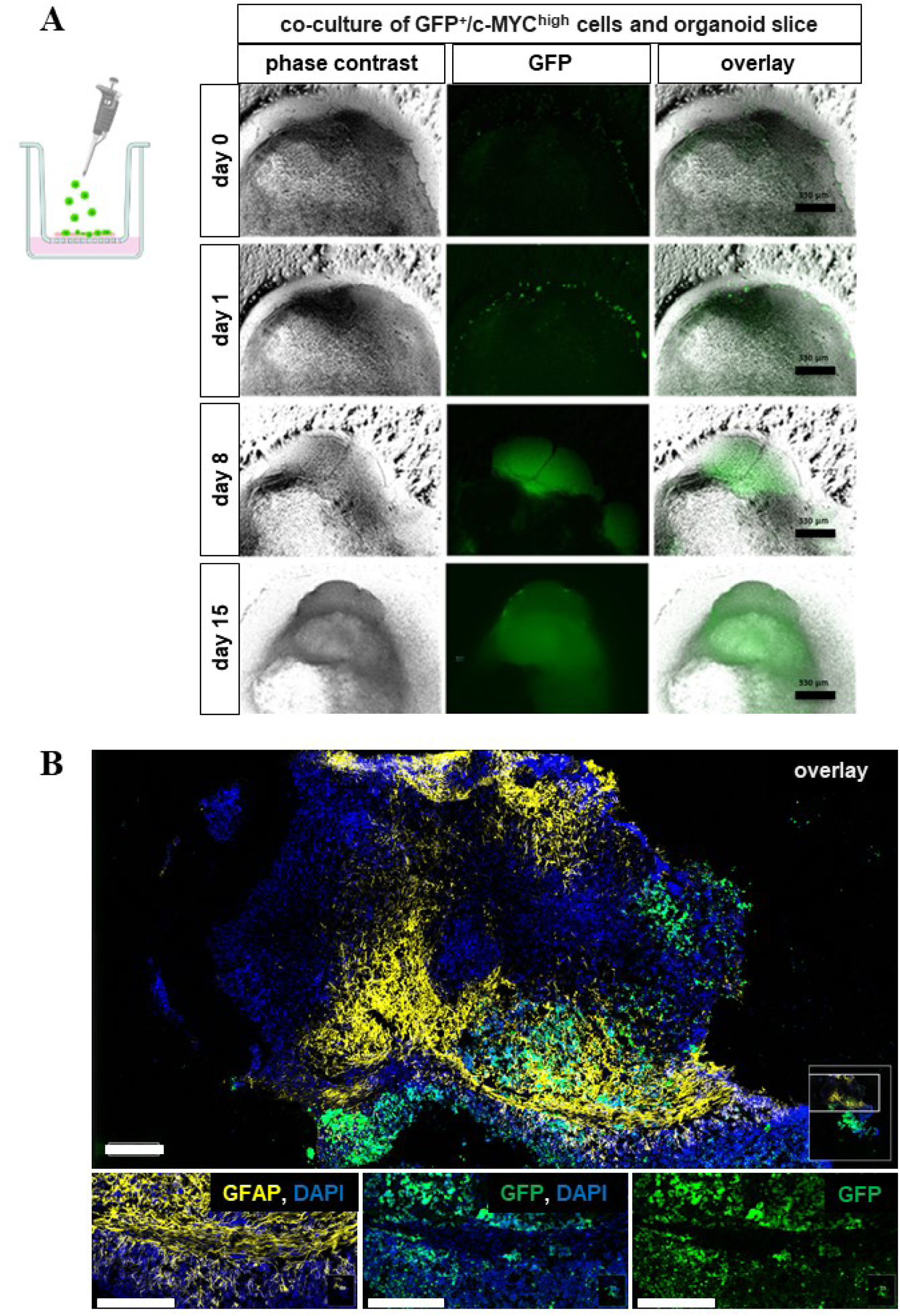
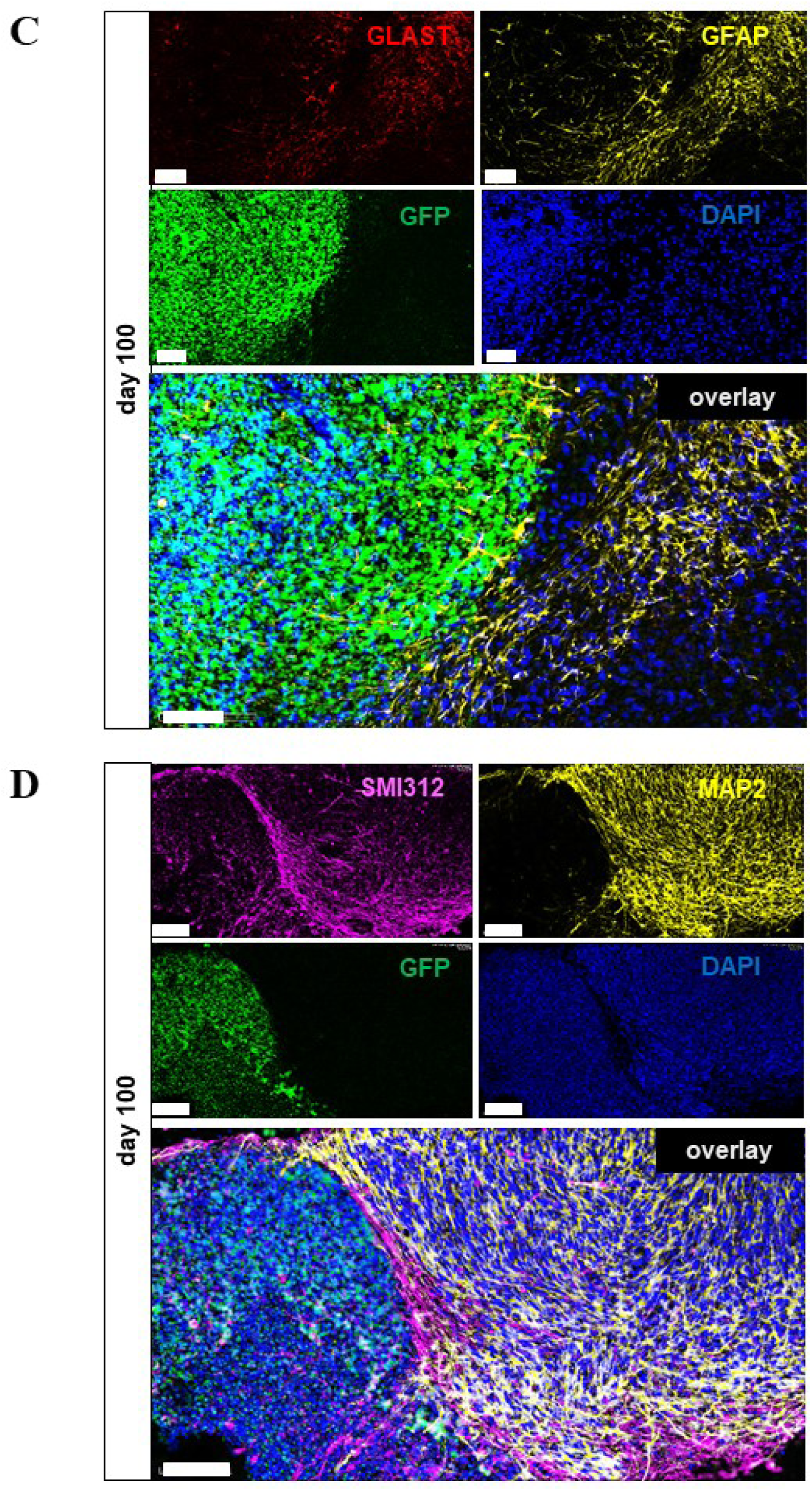
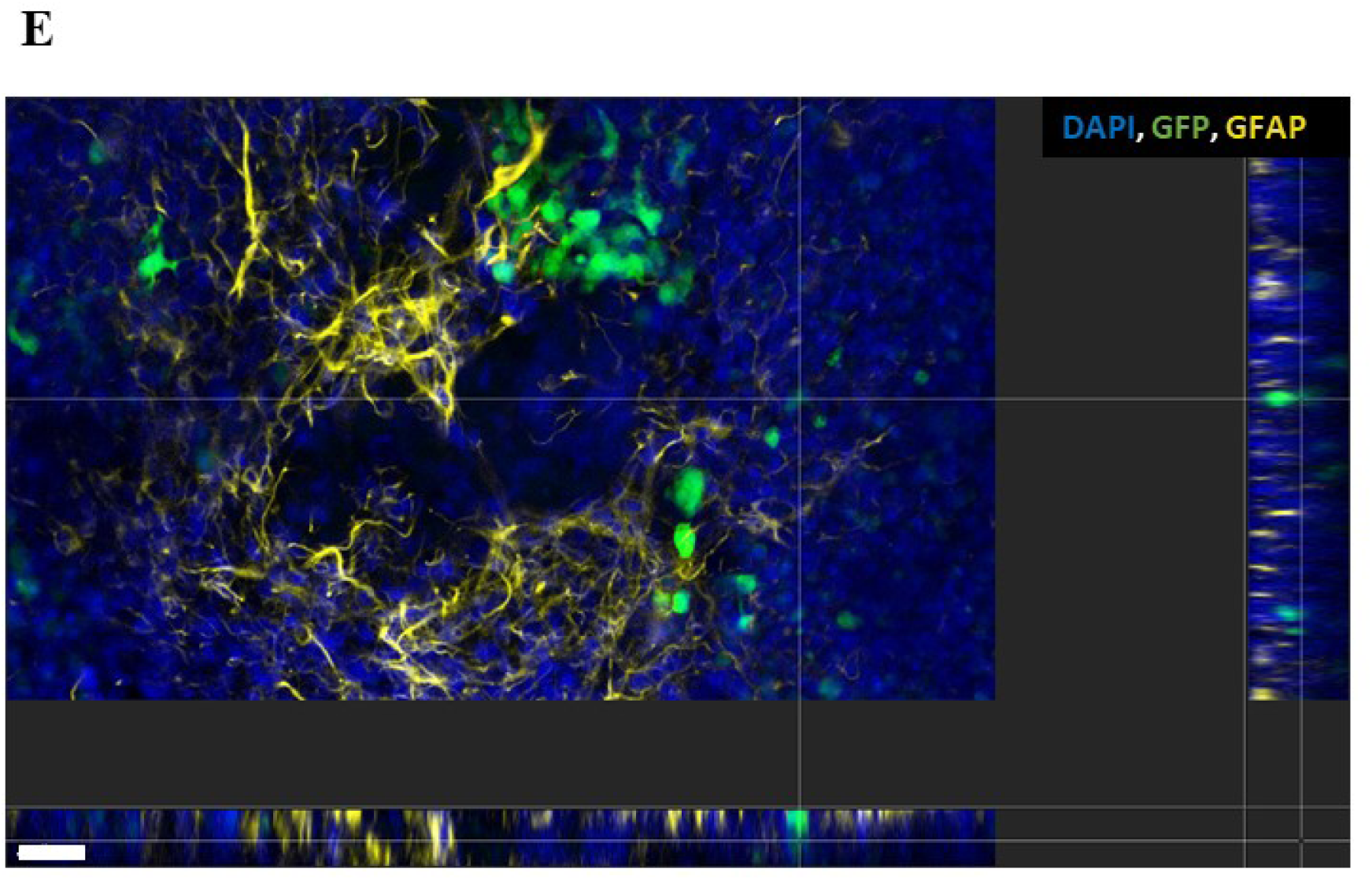
Genetically modified tumor-like cells cultured with organoids slices maintained at the air-liquid interface. **A)** Single tumor-like cells added on top of the organoid slice at ∼ day 50 of the culture show an increase in the GFP signal in time observed on 0, day 1, day 8, and day 15 of the co-culture. Scale bar: 330 µm. **B)** Representative immunofluorescence staining of GFAP (yellow) for the astrocytes of the organoid slice in the interaction with GFP^+^ cells at day 100. Scale bar: 300 µm. **C)** Representative immunofluorescence staining of GLAST (red) and GFAP (yellow) in the organoid slice containing tumor-like GFP^+^ (green) cells. Scale bar: 100 µm. **D)** Representative immunofluorescence staining of SMI312 (magenta) and MAP2 (yellow) in the organoid slice containing tumor-like GFP^+^ (green) cells. Scale bar: 30 µm. **E)** Representative immunofluorescence staining of GFAP (yellow) for the astrocytes of the organoid slice in the interaction with GFP^+^ cells (green) in the 3D image of 30 μm thick cryosection generated using z-stack. Scale bar: 30 µm.

**Figure 7.**
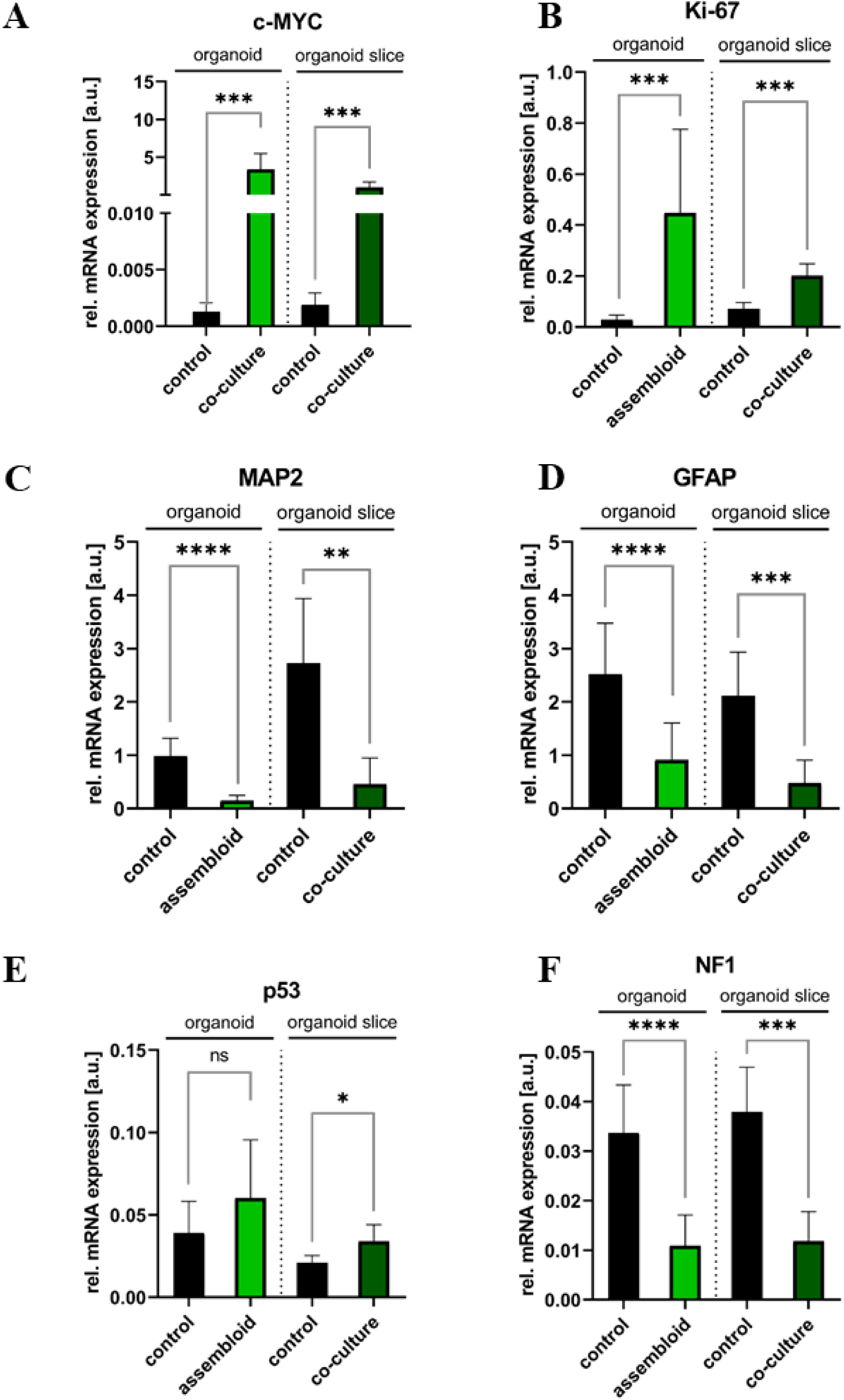
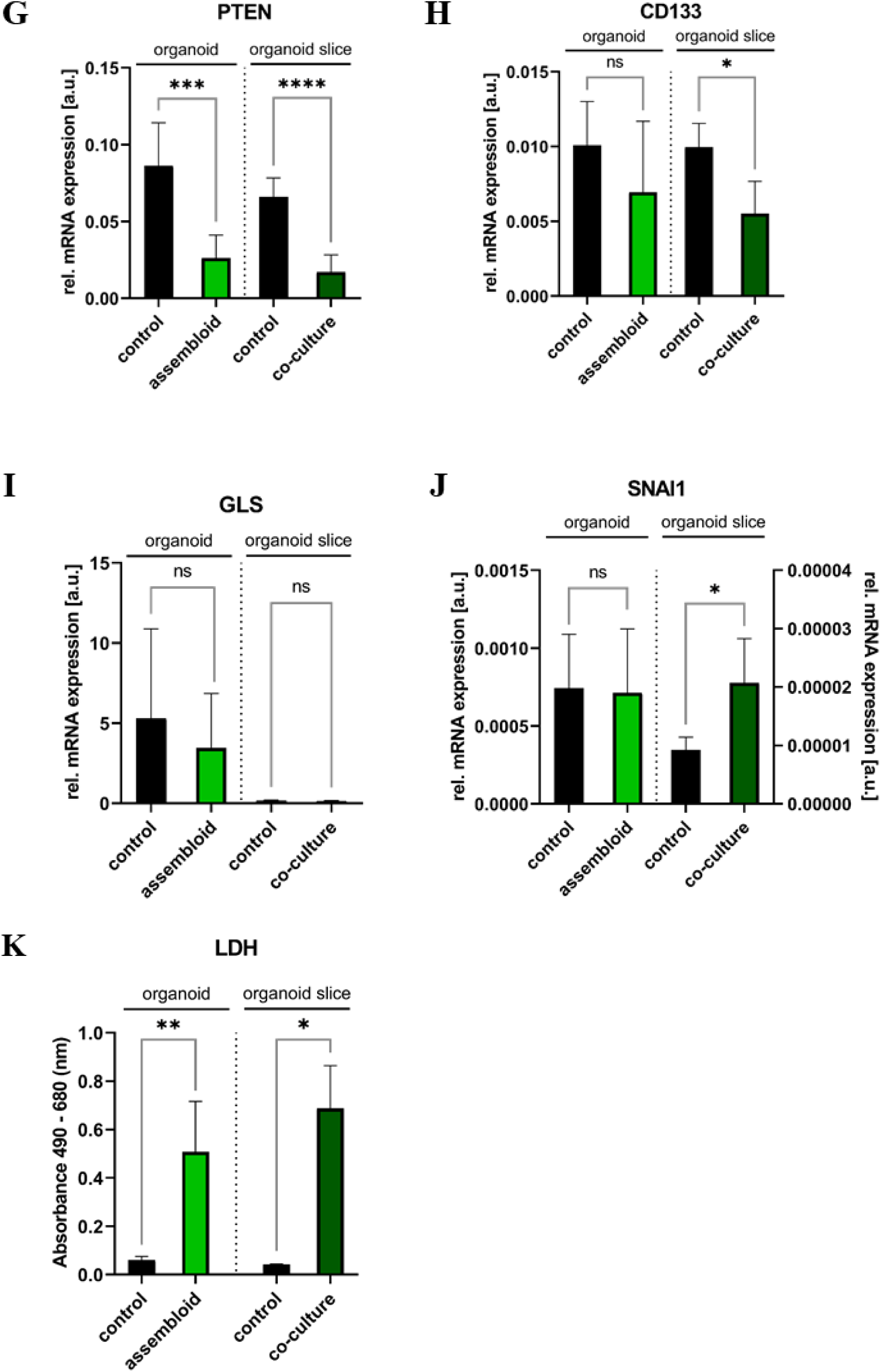
Tumor-like properties in whole organoids assembled with genetically modified tumor-like cells or in co-culture of organoid slices and tumor-like cells. A-J) Relative mRNA expression of c-MYC, Ki-67, MAP2, GFAP, p53, NF1, PTEN, CD133, GLS, and SNA-1 in the assembled whole organoids or co-cultured organoid slices with tumor-like cells compared to their respective controls. Data are presented as mean ± SD for three to six independent experiments (N = 3-6) and two to three organoids per experiment (n = 2-3), * p<0.05, ** p<0.01, *** p<0.001, **** p<0.0001. Statistical analysis was done using one-way organoids or organoid slices compared to their respective controls. Data are presented as mean + SD for three to six independent experiments (N = 3-6) and 3 to 15 organoids per experiment (n = 3-15), * p<0.05, ** p<0.01, *** p<0.001, **** p<0.0001. Statistical analysis was done using unpaired t-test.

## Discussion

The field of organoid research is advancing rapidly, with a growing body of protocols and data enabling continuous improvement of model systems for studies of organ development and disease. Human cerebral organoids, for instance, have been instrumental in advancing research on brain development, neurological and neurodegenerative diseases, and cancer [17, 26], while circumventing challenges related to the availability of human tissue and reducing the reliance on animal models. The involvement of various components of the microenvironment, including immune cells, fibroblasts, and stroma, in tumor progression is well documented [27]. However, the role of glia cells in brain tumors has not been extensively studied, despite their critical functions in supporting other cell types, e.g., through the activation of proMMP2 and the expression of AEG-1, both implicated in glioma invasion [28, 29].

### Combining traits of guided and unguided differentiation protocols leads to cerebral organoids with neuronal and glial characteristics suitable to be used as a basis for a tumor organoid model

We adopted a previously published protocol for the differentiation of pluripotent stem cells into cerebral organoids [20] with minor modifications. This protocol promotes the differentiation of glial cells, including oligodendrocytes and astrocytes. Organoids generated according to the Marton protocol lack spontaneous organization, and contain only few proliferation zones, which are specific for unguided protocols e.g. by Lancaster et al [19]. Nevertheless, the complex cellular and structural composition which is an important requirement for a 3D model system to be characterized as an organoid, was met as evidenced by immunofluorescence staining and qPCR analyses. In addition, spontaneous cell activity (calcium signals) was measured in organoid slices, but could not be detected in whole organoids, due to technical limitations of the experimental setup. However, the expression of pre- and postsynaptic markers, such as VAMP2 and HOMER, which are known to play key roles in synaptic signaling and plasticity [30, 31], in close proximity within whole organoids, serves as an indicator of functional connectivity. Previous studies showed that whole organoids are capable of spontaneous activity [32]. To increase and harmonize neuronal organization, we facilitated neural tube formation and specification via the use of microfilaments and Matrigel embedding as described by Lancaster et al. [19] before using guided differentiation according to Marton et al. to induce glial progenies [20]. This approach facilitated a bi-phasic differentiation process. The first phase, lasting approximately 50 days, was critical for cell proliferation, resulting in an increase in organoid size to approximately 3-4 mm in diameter, consistent with previous reports [33]. Beyond day 50, organoid growth plateaued, and this was corroborated by low number of Ki-67 positive cells observed at day 100. In the second phase, cell maturation of cells occurred, as evidenced by immunofluorescence staining for mature of neuronal and glial markers, such as SMI312, GFAP, and MBP. Notably, the organoids also retained progenitor cells, including those expressing nestin. Nestin, an intermediate filament protein typically expressed by NPCs (neural progenitor cells) is re-expressed in reactive astrocytes in the injured brain [34]. Neuronal marker MAP2 was detected in organoids at an early stage of differentiation (cells were MAP2^+^ and DCX^+^), and not only in more mature neurons as usually described for MAP2 [35]. DCX, which regulates microtubule dynamics and SMI312, marker for intermediate filaments supporting axonal structure, were also present in less mature cells and more differentiated neurons. CX43, a marker of cell junctions important for intercellular communication during neural development and in astrocytes, further validated the maturation process [36]. The shift in marker expression and the reorganization of cellular composition reflect the differentiation and maturation of the organoids. The minimal number of caspase-3 positive cells suggested that the organoid models remained viable throughout the culture period. However, as organoid size increases, supply with oxygen and nutrients is aggravated leading to a so-called necrotic core. Cell death in the organoid interior were confirmed by shrinked and fragmentated nuclei and an increased level of extracellularly released LDH [37]. To avoid this, we employed an air-liquid interface (ALI) technique for culturing organoid slices. Originally described for hippocampal slices [38] and later applied to cerebral organoids generated by unguided differentiation [22], the ALI technique improved oxygen and nutrient supply, reducing cell death and enabling culture maintenance for over one year. The enhanced nutrient and oxygen delivery resulted in better differentiation and maturation, particularly evidenced by the development of robust axonal networks (SMI312 staining) extending from organoid slices cultured on porous membrane inserts. These axonal tracts, which resembled subcortical projection tracts [19, 22], were more pronounced and organized than those observed in whole organoids. While astrocytic markers such as GLAST, CX43 and GFAP showed similar expression patterns in both organoids and organoid slices, lower mRNA expression of the immature astrocyte marker VIM and higher expression of the neural marker (MAP2) in organoid slices indicated a more advanced maturation of neural cells. Additionally, PDGFRα signaling, known to regulate oligodendrocyte precursor cell numbers [39], was more prominently expressed in organoid slices compared to whole organoids. This further corroborates the advanced differentiation of oligodendrocyte precursors and the presence of MBP, although at low levels in both mRNA and protein analyses, indicating that myelination at day 100 of culture is still in its early stages. In conclusion, the generated cerebral organoid models faithfully recapitulate key features of brain development, including the complex cellular composition of the cortex. With a high abundance of astrocytes, these models provide a solid foundation for investigating the role of glial cells in brain tumors, enabling closer examination of their involvement in tumorigenesis.

### c-MYC oncogene amplification as a prerequisite for the tumor organoid model

Much effort has been employed to establish *in vitro* tumor models, particularly for brain tumors, with most culture methods aiming to preserve the characteristics of the parental tumor (reviewed in Ledur et al. [40]) albeit with varying outcomes and limited scalability [41, 42]. The advent of organoid technologies has enabled the co-[43, 44]. However, this approach raises concerns regarding the mixing of distinct genetic backgrounds and developmental stages, as adult tumor tissue is combined with early embryonic cerebral organoids. This issue has been addressed through clonal mutagenesis, specifically the amplification of oncogenes and the suppression of tumor-suppressor genes, to induce tumors in brain organoids [17, 18]. In this study, we used a modified version of one of such protocols, where the c-MYC oncogene was overexpressed in cerebral organoids using the Sleeping Beauty Transposon system [18]. Given that nucleofection of cells in organoids is a random process [18] resulting in a heterogeneous distribution of genetically modified cells, we addressed this variability by isolating GFP^+^/c-MYC^high^ cells from nucleofected organoids via FACS. These isolated cells could be passaged and/or cryopreserved, enabling high-throughput scaling and usage at a self-determined time. Furthermore, these genetically modified cells were cultured in aggregates and fused to sister organoids or co-cultured with organoid slices, enhancing the reproducibility of the model and ensuring a consistent genetic background across tumor-like and normal cells. The specificity of co-expression of GFP and c-MYC were confirmed by immunofluorescence (IF) staining. GFP^+^/c-MYC^high^ cells displayed tumor-like properties, including high proliferation (Ki-67 expression), an immature and stem cell-like phenotype (SOX and CD133 expression), and invasive potential (VIM expression) in in 2D culture. Axonal marker SMI312 was detected in some cells, while glial progenitor marker PDFGRα was scarcely expressed, suggesting a neuronal rather than to glia lineage commitment for these cells. Additionally, the transcriptional profile of GFP^+^/c-MYC^high^ cells differed significantly from that of control cells in sister organoids, which lacked detectable c-MYC expression. Notably, GFP^+^/c-MYC^high^ cells exhibited elevated expression of p53, GLS and SNAI1 compared to cells of control organoids, suggesting altered bioenergetics and mesenchymal features commonly associated with tumorigenesis [45, 46], while retaining p53 regulatory mechanisms. This heterogeneity of c-MYC expression within the cell population and between batches mirrors the variability observed in human tumors [47], highlighting the challenges of tumor modeling. In nucleofected organoids, where normal cells still predominate, this variability could result in no significant changes or decreased expression of p53, Ki-67, CD133, GLS, and SNAI1 when compared to GFP^+^/c-MYC^high^ cells. As previously described, p53 is a well-known transcriptional target of c-MYC, with both c-MYC and E1A shown to stabilize p53 and trigger p53-dependent transcription [48, 49]. In our system, the expression of c-MYC was positively correlated with p53 and Ki-67 expression, consistent with its role as a prognostic biomarker in brain tumors such as gliomas and glioblastomas [50, 51]. As anticipated, the expression of tumor suppressor genes NF1 and PTEN was reduced in nucleofected organoids compared to control organoids. However, no difference in NF1 and PTEN expression between GFP^+^/c-MYC^high^ cells and control organoids suggests that these genes may be regulated by factors other than c-MYC in our model system [52, 53]. It has been reported that PTEN can acquire a pro-tumoral role by stabilizing gain-of-function p53 mutants in glioma cells [54–56]. Furthermore, CD133, a marker associated with tumor stemness and glioblastoma progression, showed a positive correlation with Ki-67 in some tumor models [57]. In our study, CD133 expression was lower in tumor-like cells compared to controls, which could be attributed to factors such as autophagy or lactate levels in the tumor microenvironment [58, 59].

The tumor-like properties of GFP^+^/c-MYC^high^ cells were further confirmed in our tumor models, where the GFP signal increased over time and GFP^+^ cells were detected in organoids or organoid slices, reflecting the proliferative and invasive potential of these cells. c-MYC overexpression has been shown to contribute significantly to the aggressiveness of medulloblastoma [60, 61], although the effects of exogenous c-MYC expression via the Sleeping Beauty system may differ from endogenous c-MYC in patient tumors. Mesenchymal features of the tumor model were corroborated by the expression of SNAI1 [45]. No significant changes in GLS expression between tumor models and controls suggest that glutamine metabolism, a key feature in tumor progression, remains unaltered in our system [62]. Additionally, an increase in extracellular LDH levels in co-cultures may be associated with the massive proliferation and overgrowth of GFP^+^/c-MYC^high^ cells and the associated increased competition for space and nutrients, a hallmark of many brain tumors [63, 64]. The gene expression profiles obtained from nucleofected whole organoids and co-cultured systems were highly consistent, demonstrating the robustness of the co-culture model in providing more reliable and physiologically relevant results compared to traditional 2D cultures. Moreover, this approach allows for experiments at defined and later time points, including long-term culture, which is not feasible with random nucleofection without FACS sorting.

Genetically modified GFP^+^/c-MYC^high^ cells in spheres benefited from the cultivation with normal organoids, exhibiting increased cell density compared to spheres alone. Normal organoids provided not only a larger surface for tumor-like cells to spread but also extended GFAP-positive astrocytic protrusions into the sphere. This supports the known role of astrocytes and oligodendrocytes in tumor progression and invasion through secretion of various factors [65, 66]. Tumor cells can also activate astrocytes via the JAK/STAT pathway, promoting an immunosuppressive environment [67]. Interestingly, while neuronal cells expressing axonal markers (SMI312) and MAP2 were present in our model, they were spatially segregated from GFP^+^ cells and appeared less permissive to tumor cell invasion, a finding consistent with xenograft models where glioblastoma cells predominantly invade astrocyte- and microglia-rich regions rather than neuronal areas [68]. Despite this, a small proportion of GFP^+^/c-MYC^high^ cells were able to migrate into normal organoid tissue or infiltrate organoid slices, resembling the behavior of disseminated tumor cells in glioblastomas [69].

While GFAP, a marker of astrocytomas, showed minimal expression in highly proliferative GFP^+^ cells, it does not correlate directly with malignancy grade, as different GFAP isoforms can coexist in these tumors [70]. However, c-MYC overexpression alone is insufficient to induce a glioblastoma-specific phenotype. Future improvements to this model will involve introducing deletions in tumor suppressor genes commonly affected in pediatric glioblastoma and conducting single-cell analyses to further refine the model.

## Conclusion

The 3D microenvironment of organoids or organoid slices, which exhibit key features of the brain, including a high proportion of glial cells - particularly astrocytes - provides a supportive framework for the proliferation, migration, and invasion of genetically modified cells with tumor-like characteristics. We have developed a robust and reproducible model system utilizing two distinct culture approaches, enabling the long-term study of various treatment modalities, such as radiotherapy, chemotherapy, and immunotherapy, in an autologous context. This approach ensures that all cells originate from the same genetic background, offering a more accurate and personalized platform for therapeutic evaluation.

## Methods

### Human embryonic stem cell culture

The feeder-independent hES cell line WA09-FI (H9) was used for all studies presented as approved according to §4 and §6 of the German Stem Cell Act (registry numbers 3.04.02/0125 and 3.04.02/0125-E01). The line was originally generated by the group of Dr. James Thomson at the University of Wisconsin [71]. H9 cells were obtained from the WiCell Research Institute, Wisconsin, USA, at passage 23 and were used for experiments in passages 34-56. Cells were routinely cultured on Laminin-521-coated culture dishes (BioLamina, #600962, 10 µg/ml) in mTeSR^™^1 basal medium medium with addition of 1x mTESR^™^1 supplement (STEMCELL Technologies) and 50 U/ml penicillin and 50 µg/ml streptomycin (Merck, #A2212) or in TESR^™^-E8^™^ medium with 1xTesr^™^-E8^™^ supplement. Cells were passaged every week using ReleSR (STEMCELL Technologies, #05872). Briefly, medium was aspirated, and cells were rinsed first with 2 ml PBS without calcium chloride and magnesium chloride (PBS-/-) and then with 200 µl ReleSR. After aspiration of the non-enzymatic passaging reagent, cells were incubated for 2 min at 37°C to detach only pluripotent cells. Detachment was stopped by addition of pre-warmed medium, and cells were seeded to about 1.0 x10^5^ cells per 60 mm Petri dish.

### Plasmid constructs

Overexpression of c-MYC oncogene in cells of organoids was based on the Sleeping Beauty transposase system consisting of 2 vectors: SB100X (VB190927-1101xtm) and c-MYC (VB190927-1100geg) expressing vector (VectorBuilder).

### Generation of cerebral organoids

Cerebral organoids were generated as previously described [19, 20]. Briefly, for the generation of embryonic bodies (EBs), H9 cells were detached using ReLeSR (Stemcell Technologies) for 3 min at 37 °C. 18000 cells in TeSR^™^-E8^™^ medium with addition of 50 µM Rho-associated protein kinase (ROCK) inhibitor (Tocris Bioscience), to enhance the survival of the dissociated cells by preventing apoptosis and to improve EB formation, were plated into each well of an U-bottom suspension plate (Sarstedt) pre-coated with anti-adherence solution (Stemcell Technologies) for 5 min, 1000 rpm, at RT. Microfilaments (surgical suture) or 0.4% polyvinyl alcohol (PVA) were used to enhance the assembly of the embryoid body to give rise to the organoid [72, 73]. Medium was changed to Essential 6^™^ Medium (Thermo Fisher) supplemented with two SMAD pathway inhibitors named dorsomorphin (2.5 μM, Sigma) and SB-431542 (10 μM, R&D Systems) two days after seeding. This day is defined as the start of differentiation and therefore, the age of the organoids was calculated from this day (0 days old; d0). For the first 5 days, Essential 6^™^ medium was changed every day and supplemented with dorsomorphin and SB-431542 to derive neural progenitor cells from hES cells. On day 6, the organoids were transferred to 24-well plates and cultured in an improved differentiation medium – A (IDM-A, according to Lancaster et al., 2017) containing 1:1 Dulbecco’s Modified Eagle Medium/Nutrient Mixture F-12 (DMEM/F12) and Neurobasal^®^, 0.5% N-2 supplement, 2% B27^®^ – vitamin A (Thermo Fisher), 0.25% insulin, 50 μM β-mercaptoethanol, 1% GlutaMAX^™^-1, 0.5% Minimum Essential Medium – Non-essential amino acid solution (MEM-NEAA), and 1% penicillin-streptomycin (Sigma). The IDM-A was supplemented with 20 ng/mL epidermal growth factor (EGF, Peprotech) and 20 ng/mL basic fibroblast growth factor (bFGF, Peprotech) for 19 days (until day 24) with daily medium change in the first 10 days, and every other day for the subsequent 9 days, to promote neural and glial lineage differentiation. Additionally, 5 μM of the inhibitor of WNT Production-2 (IWP-2, Selleckchem) was added to the medium from day 4 until day 24 to inhibit the WNT pathway, and 1 μM of the small molecule smoothened agonist (SAG, Sigma) was added from day 12 to day 24 to activate the sonic hedgehog (SHH) pathway, to promote further neuronal proliferation and maturation. EBs were embedded into Matrigel (Corning) droplets on day 15 and cultured in IDM-A with addition of EGF/bFGF, IWP2 and SAG with medium being exchanged every second day. From day 25-36, organoids were cultured in IDM-A supplemented with 60 ng/mL Triiodothyronine (T3, Sigma), 100 µg/mL biotin (Sigma), 20 ng/mL neurotrophin-3 (NT-3, Preprotech), 20 ng/mL brain-derived neurotrophic factor (BDNF, Peprotech), 10 μM cyclic adenosine monophosphate (cAMP, Sigma), 5 ng/mL hepatocyte growth factor (HGF, Peprotech), 10 ng/mL insulin-like growth factor 1 (IGF-1, VWR), and 10 ng/mL platelet-derived growth factor AA (PDGF-AA, R&D Systems), to promote OPC survival and proliferation. To allow homogeneous distribution of the different factors and nutrients, on day 25, the organoids were transferred to T25 suspension flasks with filtered cap and put on an orbital shaker with a throw of 19 mm and a corresponding speed of 62 r.p.m. From day 37 onwards, organoids were cultured in IDM+A, namely IDM-A containing 2% B27^®^ + vitamin A, plus 0.4 mM vitamin C and 12.5 mM HEPES Buffer Solution to control pH levels. The medium was again supplemented with T3, biotin and cAMP to promote oligodendrocyte maturation. Media changes were performed every three days until day 49, then every three to four days for the duration of the culture.

### Induction of genetic modifications in organoids

Cerebral organoids were subjected to nucleofection on day 11 of the culture using Amaxa™ P3 Primary Cell 4D Nucleofector X Kit (Lonza) according to the manufacturer’s protocol. Briefly, 8-15 organoids were placed between electrodes of the cuvette and the excess medium was removed. Then, nucleofection medium consisting of premixed solution (82 µl) and supplement (18 µl) from the kit with addition of plasmid vectors, SB100X (12.8 µg) and c-MYC (1.1 µg or 2.2 µg) expressing vector (VectorBuilder), was added to organoids. Sham-nucleofected organoids served as a negative control. Nucleofection program CB-150 for H9 cells was applied and organoids were immediately placed at 37 °C and 5% CO_2_ for 10 min, following addition of culture medium to the cuvette and incubation for the next 10 minutes. Then, organoids were placed in a Petri dish and maintained in IDM-A medium with addition of EGF, bFGF and IWP-2. Medium was exchanged according to the same protocol as described for normal control organoids in a previous section. Nucleofected organoids were cultured in parallel with control sister organoids (i.e. from the same batch). Organoids were monitored daily for the presence of GFP signal which was associated with successful nucleofection.

### Isolation of genetically modified cells from cerebral organoids for co-culture with sister organoids

Nucleofected organoids (2-8 per sample) showing large areas with GFP signal were selected for isolation of genetically modified cells at day 50-91 of culture. Organoids were then washed in PBS-/- and dissociated by mincing and using StemPro^®^ Accutase^®^ in a 15-30 min incubation in a microtube at 37 °C and 5% CO_2_ on an orbital shaker. Additionally, every 10 min the suspension was pipetted up and down and the supernatant containing single dissociated cells was transferred to a new microtube and centrifuged at 300xg for 5 min or 200xg for 7 min at RT. The cell pellet was resuspended in PBS-/-with addition of 50 µM Rock inhibitor (Tocris Bioscience) and placed on ice to minimize the cell death. The remaining non-dissociated organoid pieces were used to gain more cells by repeating the incubation with new addition of StemPro^®^ Accutase^®^, centrifugation at 300xg for 5 min or 200xg for 7 min at RT. Cell suspension from the same condition was merged after dissociation and filtered using a cell strainer with 35 µm nylon mesh (Corning) into a cell sorting tube. Cells were immediately sorted by fluorescence-activated cell sorting (FACS) based on the presence of GFP signal. Sham-nucleofected or control organoids dissociated using the same procedure were used as a negative control for adjustment of gates for cell populations prior to sorting. Positively selected (GFP^+^) cells were plated onto Geltrex™-coated petri dishes and cultivated in IDM+A supplied with 60 ng/ml T3, 100 µg/ml biotin and 10 μM cAMP for their propagation and used for generation of spheres (1×10^4^ cells mixed with 5 µl Matrigel), namely tumor spheres, to be assembled with whole organoid. For this, one organoid and one tumor sphere were cultivated together in one well of a 24-well suspension plate at an angle on an orbital shaker with 19 mm throw at 114 r.p.m. The exchange of IDM+A was performed every two days, while the medium was supplied with T3, biotin and cAMP. Alternatively, GFP^+^ cells were applied directly after sorting on top of the slice at ALI-culture (1×10^3^ cells/organoid slice).

### Air-liquid interface culture

Culture of organoids at air-liquid interface was performed according to the protocol by Giandomenco and colleagues [20] with slight modifications. Briefly, 3-6 cerebral organoids between 50 and 60 days old were collected and washed in Hanks’ Balanced Salt Solution without Ca^2+^ and Mg^2+^ (HBSS, Thermo Fisher) and embedded in 3% low-melt agarose (Sigma) at approximately 37 °C in peel-a-way® embedding molds (Sigma). The agarose blocks with embedded organoids were incubated on ice for 10–15 min and cut into 300 μm thick sections in cold HBSS using Leica VT1200S vibratome. An amplitude of 1 mm, a speed of 0.30 mm/s, and a razor blade angle of 21° were used. Sections were collected onto Sarstedt TC-inserts (pore size of 0.4 μm) in 12-well plates and first left to equilibrate for 1 h at 37 °C in IDM+A and addition of 100 µg/ml Biotin, 60 ng/ml T3 and 10 µM cAMP. Cultured slices were maintained at the air-liquid interface in IDM+A medium with the addition of mentioned supplements at 37 °C and 5% CO_2_ with daily medium changes (3/4 of the medium).

### Immunofluorescence

Organoids and organoid slices were fixed in 3.7% formaldehyde (Carl Roth) at 4 °C overnight and then washed three times for 5 min with PBS. Organoids were dehydrated in sucrose gradient (7%, 10% 30%, 40% 60% sucrose in PBS; 4h for each step except overnight for 30% and 60% sucrose at 4 °C). Organoid slices were dehydrated in 30% sucrose (Sigma) in PBS overnight. Organoids were then embedded in 7.5% gelatin (Neolab)/10% sucrose using in house 3D-printed PDMS embedding molds. Embedded organoids were frozen on dry ice and stored at −80 °C prior cryosectioning at 10 µm (if not stated differently) using CM1860 cryostat (Leica Biosystems). For staining against MBP, antigen retrieval was performed using Tris-EDTA (pH 9) for 30 min in water bath at 95 °C. Cryosections were blocked and permeabilized in 0.5% Triton X-100 (ThermoFisher Scientific)/1% BSA (Carl Roth) in PBS for 30 min and blocked with 1% BSA in PBS for 30 min at room temperature (RT). Incubation of samples with primary antibodies in 1% BSA in PBS was performed either for 1 h at RT or at 4 °C overnight following washing three times for 5 min with PBS. If the primary antibody was not directly conjugated to a fluorescent dye, samples were then incubated with secondary antibodies in 1% BSA in PBS for 1h at RT followed by washing three times for 5 min with PBS and incubation with 5 µg/ml DAPI for 4 min for nuclei staining. Then, the samples were washed two times for 5 min in PBS followed by washing for 5 min with Millipore water prior to mounting with fluorescence mounting medium (Dako). The primary and the secondary antibodies used for immunofluorescence are listed in Supplementary Tables 1-3. Images were captured with a fluorescence microscope Zeiss Axio Imager.Z2 equipped with Metafer5 software (Metasystems) and with a confocal microscope DMI 4000B (Leica Microsytems) with LAS X software (v3.5.7.23225, Leica Microsystems). The stainings were performed on samples from three independent preparations with at least two organoids per group. Images were processed with ImageJ (v1.53i, National Institute of Health (NIH)).

### Clearing and immunofluorescence staining for light sheet microscopy

Clearing, staining and imaging of whole organoids were performed according to the modifications of the CUBIC protocol [74–76]. Fixed, whole organoids were treated with 50 % CUBIC-L (TCI, #T3740) in Millipore overnight at 37 °C on a shaker set to 144 rpm. The organoids were then incubated in CUBIC-L at 37 °C for 2-3 days with agitation at the same speed, followed by three washing steps with PBS, 1 h each at RT. Subsequent incubations were performed at RT with gentle agitation (60 rpm), taking care to avoid damage of organoids during solution changes. For immunofluorescence staining, the organoids were blocked with 5% BSA in PBS for 3 h at 37 °C and incubated with primary antibodies diluted in 5% BSA in PBS for 24 h. The following day, organoids were washed three times with PBS (1 hour each) and incubated with the corresponding secondary antibodies in 5% BSA in PBS for 24 h. Afterwards, unbound antibodies were removed with three 1-hour washes in PBS. If applicable, conjugated antibodies diluted in 5% BSA in PBS, were applied and incubated for 24 h. For the final step, the last antibody solution was washed out with three 1-hour PBS washes. The organoids were then stained with DAPI (5 µg/ml) for at least 12 hours and were washed three times with PBS (1 hour each) afterwards. To match the refractive index, the organoids were first treated with 50% CUBIC-R(M) (TCI, #T3741) in Millipore for 12 hours, followed by incubation in CUBIC-R for 2 days, with the solution refreshed after 24 hours. Cleared and stained organoids were imaged using a Leica STELLARIS DLS system with an HC APO L 10x/0.30 W DLS objective and a 7.8 mm TwinFlect mirror cap. Samples were mounted in a 35 mm glass-bottom dish (ibidi), with a base layer of 4% agarose/CUBIC-R poured and set for 30 minutes. The organoids were then embedded on top of this layer in the same matrix. Areas for the mirror cap were cut out, and the dish was filled with CUBIC-R solution before immersion of the objective. Images were processed with ImageJ (v1.53i, National Institute of Health (NIH)) and Huygens™ Essential (Scientific Volume Imaging).

### Real time RT-PCR analysis

C-MYC^high^ cells, tumor spheres (n = 4-6 per batch), whole organoids (n = 3 per group) or organoid slices (n = 3 per group) were collected separately or pooled in QIAzol Lysis Reagent (Qiagen, #79306) and total RNA was isolated using the Qiagen RNeasy Mini Kit (#74106) according to the manufacturer’s instructions including a DNA removal step using the RNase-free DNase Set (Qiagen, #79254). 50 ng RNA were reverse-transcribed via the RevertAid RT Kit (Life Technologies, #K1691). Relative RNA expression was analyzed using the Hot FIREPol EvaGreen qPCR Mix Plus from Solis Biodyne (08-24-0000S) and the QunatStudio 3 Real-Time PCR System. The standard curve for each primer was generated using either human fetal and or adult brain mRNA. Target expression level of each gene was normalized to 18S rRNA. Primer sequences are listed in Supplementary Table 4.

### Lactate dehydrogenase (LDH) level

Extracellular lactate dehydrogenase (LDH) released from necrotic cells in the media was determined using CyQUANT™ LDH Cytotoxicity Assay Kit (Invitrogen) according to manufacturer’s instructions. Briefly, culture medium was collected at approximately day 100 of the culture and stored at −80 °C. On the day of assay, 50 µl of each sample medium was added to a 96-well black clear-bottom plate in duplicate wells. Then, 50 µl of Reaction Mixture from the kit was added to each sample and incubated at room temperature for 30 minutes in the dark to enable conversion of lactate to pyruvate in a reaction catalyzed by LDH where NAD+ is reduced to NADH. The reaction was stopped by adding 50 µl Stop Solution from the kit. The absorbance of red formazan product generated by reduction of tetrazolium by diaphorase, which also oxidizes NADH, was measured spectrophotometrically at 490 nm. 680-nm absorbance value (background) was subtracted from the 490-nm absorbance. Obtained absorbance values are directly proportional to the amount of LDH released into the media.

### Functional analyses: Ca-imaging

The samples were cultured in the air-liquid interface as previously described. To avoid damaging the outgrowths of the samples, the slices were left on the air-liquid interface. One day before staining, the slices were covered with NB+ medium (NB+ medium, 0.5 mM glutamine, 2 % B-27+ supplement and 1 % Pen/Strep, all from ThermoFisher Scientific). The next day, the slices were loaded with a solution of 5 µM Calbryte™ 520 AM (AAT Bioquest, Inc.) in NB+ medium and incubated at 37 °C for 60 minutes. The dye solution was replaced with fresh medium to remove excess probes. Prior to recording, samples were stored in the incubation chamber of the microscope at ∼37 °C for an additional 15 minutes. The recordings were performed with a confocal inverted microscope, Leica DMI8-CS, Stellaris 5, and a multi-immersion objective HC PL APO 20x/0.75 IMM CORR CS2. The tunable Wightlight laser (WLL) was set to a wavelength of 493 nm. A maximum laser power of 15 per cent was used to protect the samples. The video sequences were recorded at 4 Hz and a size of 512 x 512 pixels. Multiple cells were segmented and analysed using the open source software ImageJ (FIJI) [77]. The fluorescence change over time was defined as ΔF/F = (F0 −F1)/(F0), where F0 represented the 10th percentile of the signal in a rolling window.

### Single nuclei RNA sequencing and data analysis

Organoid slices (n = 3) and organoids (n = 3), each pooled from the same biological replicate in one tube, were washed in cold PBS at 300xg, 5 min. After complete removal of PBS, organoid slices and organoids were placed on dry ice for 5 min, and then stored in liquid nitrogen. Preparation of samples for sequencing including the isolation of the nuclei, initial quality control, library preparation, sequencing and initial data processing were performed as a service by GENEWIZ/AZENTA (USA). The RNA-seq library was prepared using 10X Genomics Chromium^®^ 3‘ expression, and the 10X library sequencing was performed on the Illumina series. The target number of cells per sample was 6000, and number of reads per cell 50000. Raw reads were preprocessed and aligned to the human genome (hg38) using the Cell ranger pipeline (10X Genomics). Initial quality control and visualization of the data was performed in the Loupe browser (10X Genomics). Data analysis of the filtered feature matrix was performed using the R software (https://www.r-project.org/) with Seurat package and Seurat workflow (https://satijalab.org/seurat/). Cells that had unique feature counts over 2500 or less than 200, and cells that had >5% mitochondrial count were filtered out from the dataset. The data was normalized using a SCTransform function for data normalization, scaling and reduction prior to graph-based clustering and visualization of the merged and integrated datasets for organoids and organoid slices.

### Statistics

All analyses were performed for independent experiments (N) containing multiple organoids (n) and using GraphPad Prism (v 9.3.1). Normal distribution of studied variables was proved by normality and lognormality tests with accompanied QQ-plots, and homogeneity of variances was checked using F-test. Statistical comparisons were performed as stated in figure legends and included unpaired two-tailed test with or without Welch correction, Brown-Forsythe/Welch, or mixed ANOVA model either with Dunnett’s or Tukey’s post-test. * p<0.05, ** p<0.01, *** p<0.001, **** p<0.0001. Samples of organoid slices were randomly assigned to different conditions or treatments. No statistical methods were used to pre-determine sample sizes. Because of the nature of the treatment (nucleofection), data collection and analysis were not performed blind to the conditions of the experiments.

## Acknowledgements

The authors would thank members of Stem Cell Differentiation and Cytogenetics Group led by I. Schroeder for helpful discussions and valuable suggestions and particularly to Emilia Choman, Dr. Carola Hartel and Jennifer Persigehl for excellent technical assistance. We thank to Daniela Di Giovanni for her invaluable support in the establishment and validation of immunofluorescence staining protocols during her master’s thesis. This work is supported by the German Federal Ministry of Education and Research (02 NUK 049A) and the NIH grant 1RO1CA256848-01.

## Authors contribution

T.B. and E.S. planned and performed experiments, analyzed data, interpreted results and wrote the manuscript. L.K. performed confocal imaging and contributed to the interpretation of results. S.H. planned and performed Ca imaging experiments and analyzed data. M.M. and C.T. supervised electrophysiology experiments and interpreted data. I.S., D.R.G. and M.D. conceived and supervised the project and data interpretation.

