## Supplementary information for "Human cerebral organoids model tumor infiltration and migration supported by astrocytes in an autologous setting"

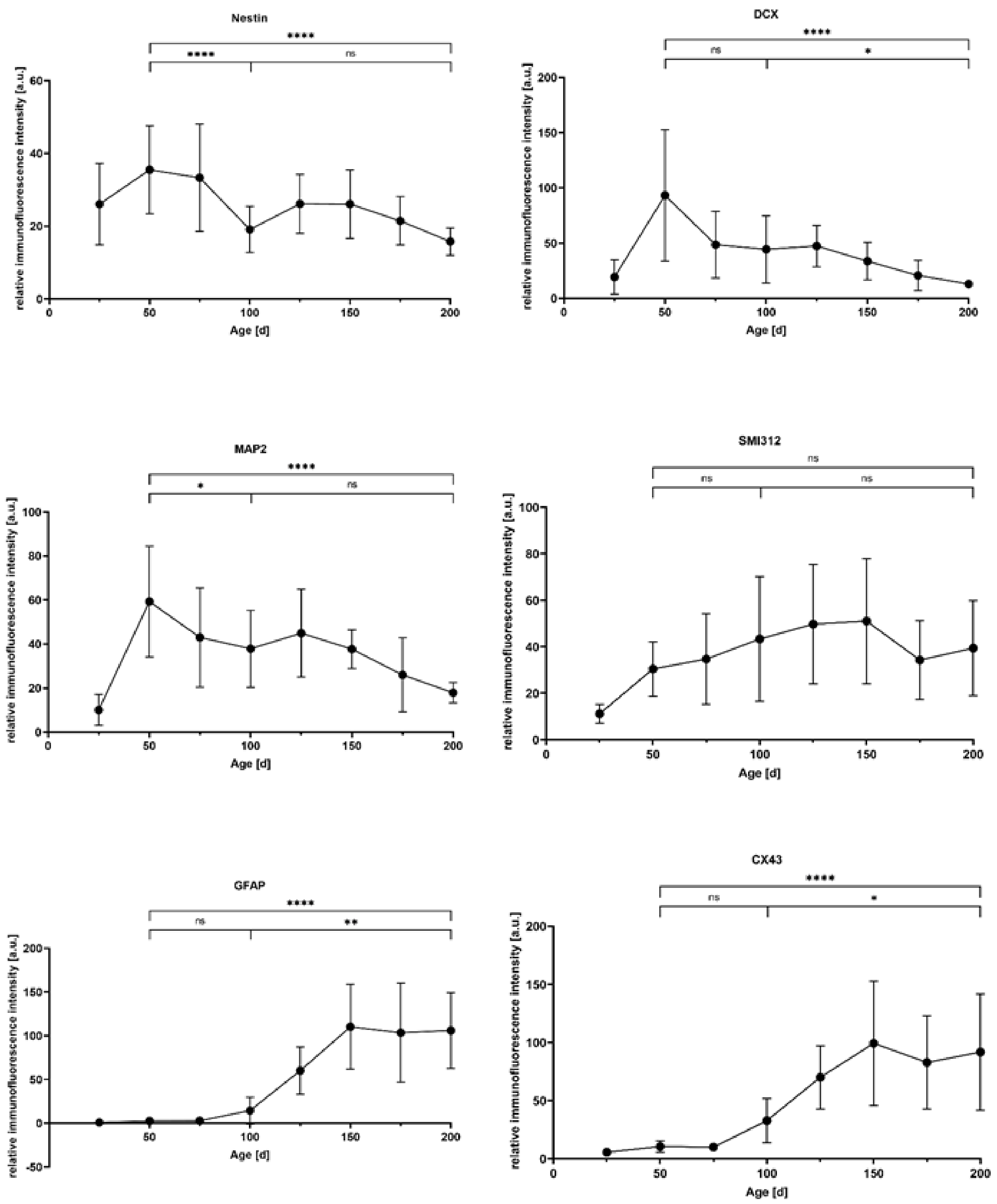


**Supplementary Figure 1. Maturation of cerebral organoids**. Relative immunofluorescence intensity of nestin, DCX, MAP2, SMI312, GFAP, and CX43 normalized to the related nuclei area. For each marker, four whole organoid sections from 25, 50, 75, 100, 125, 150, 175, and 200 days old organoids were analyzed. Data are presented as mean ± SD for two to three independent experiments (N = 2-3) and one to three organoids per experiment (n = 1-3), * $p<0.05$, ** $p<0.01$, *** $p<0.001$, **** $p<0.0001$. Statistical analysis was done using one-way ANOVA with Tukey's post-test (nestin, MAP2, and SMI312) or with Kruskal-Wallis Test with Dunn's post-test.

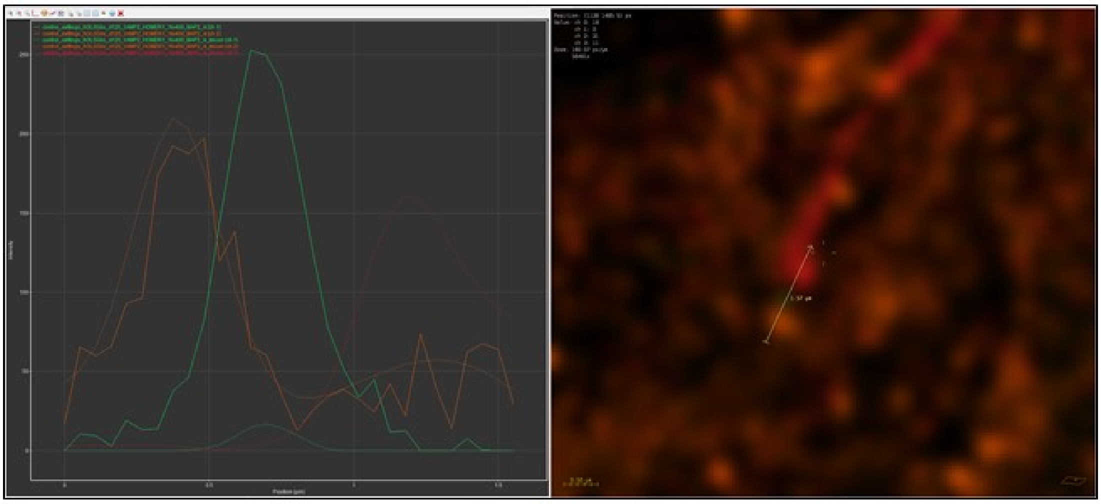

**Supplementary Figure 2. Visualization of the synaptic cleft.** Intensity measurement of fluorescent signals for presynapse (VAMP2, orange) and postsynapse (HOMER1, green) (left image). The x-axis is defined by the arrow shown in the right picture. Even with confocal imaging, the small synaptic cleft (200 to 300 nm) can be measured/visualized, especially after deconvolution (dotted line). The right picture shows the data after deconvolution. Scale bar: 0.5 µm.

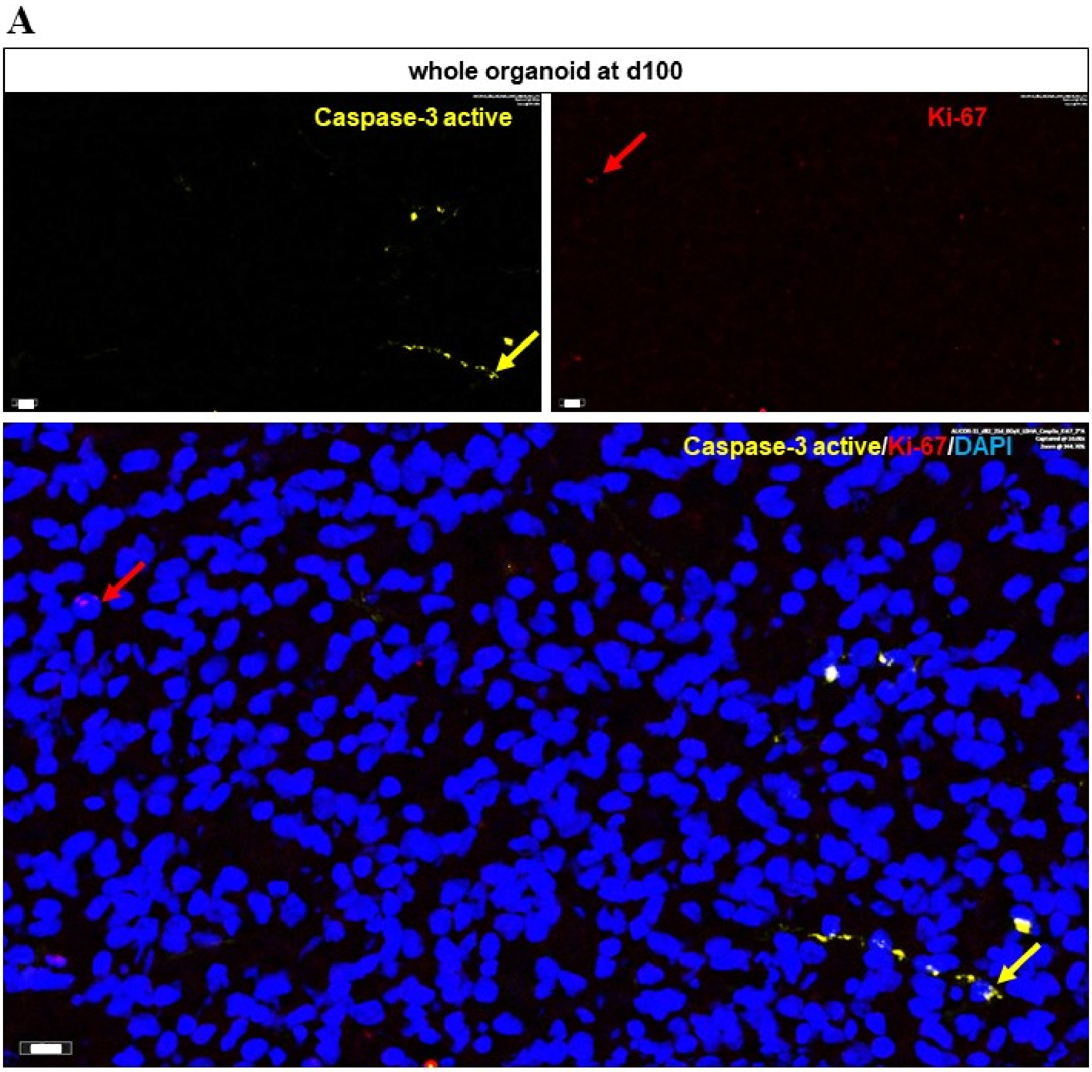

B

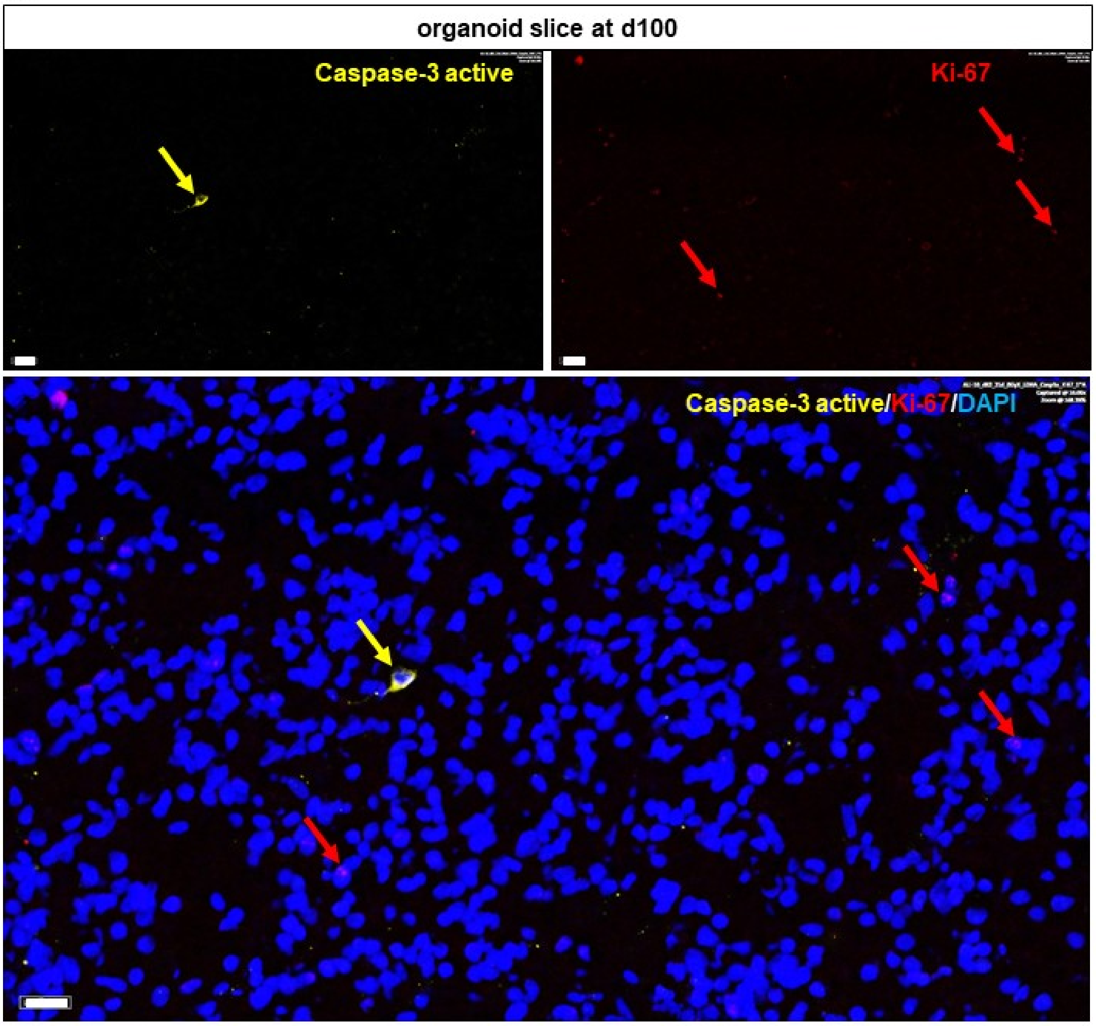


C

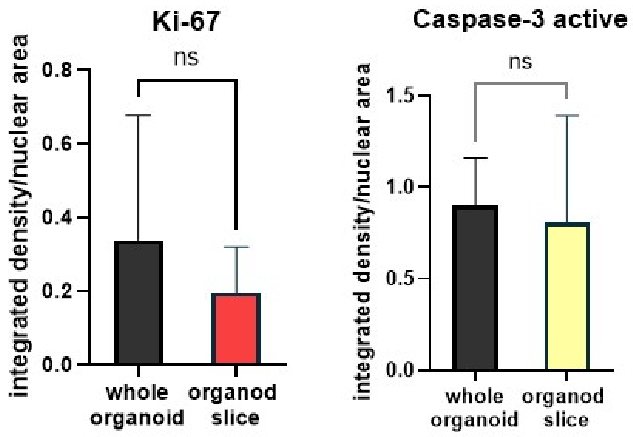


**Supplementary Figure 3. Proliferation and apoptosis in whole organoids and organoid slices. A)** Representative immunofluorescence staining of caspase-3 active (yellow) and Ki-67 (red) for 100 days old whole organoid. Scale bar: 10 µm. **B)** Representative immunofluorescence staining of caspase-3 active (yellow) and Ki-67 (red) for 100 days old organoid slice. Scale bar: 20 µm. **C)** Integrated density normalized to the related nuclei area of Ki-67 and caspase-3 active in 100 days old whole organoids and organoid slices. Data are presented as mean ± SD for three independent experiments (N = 3) and two to three organoids per experiment (n = 2-3), * p<0.05, ** p<0.01, *** p<0.001, **** p<0.0001. Statistical analysis was done using unpaired t-test.

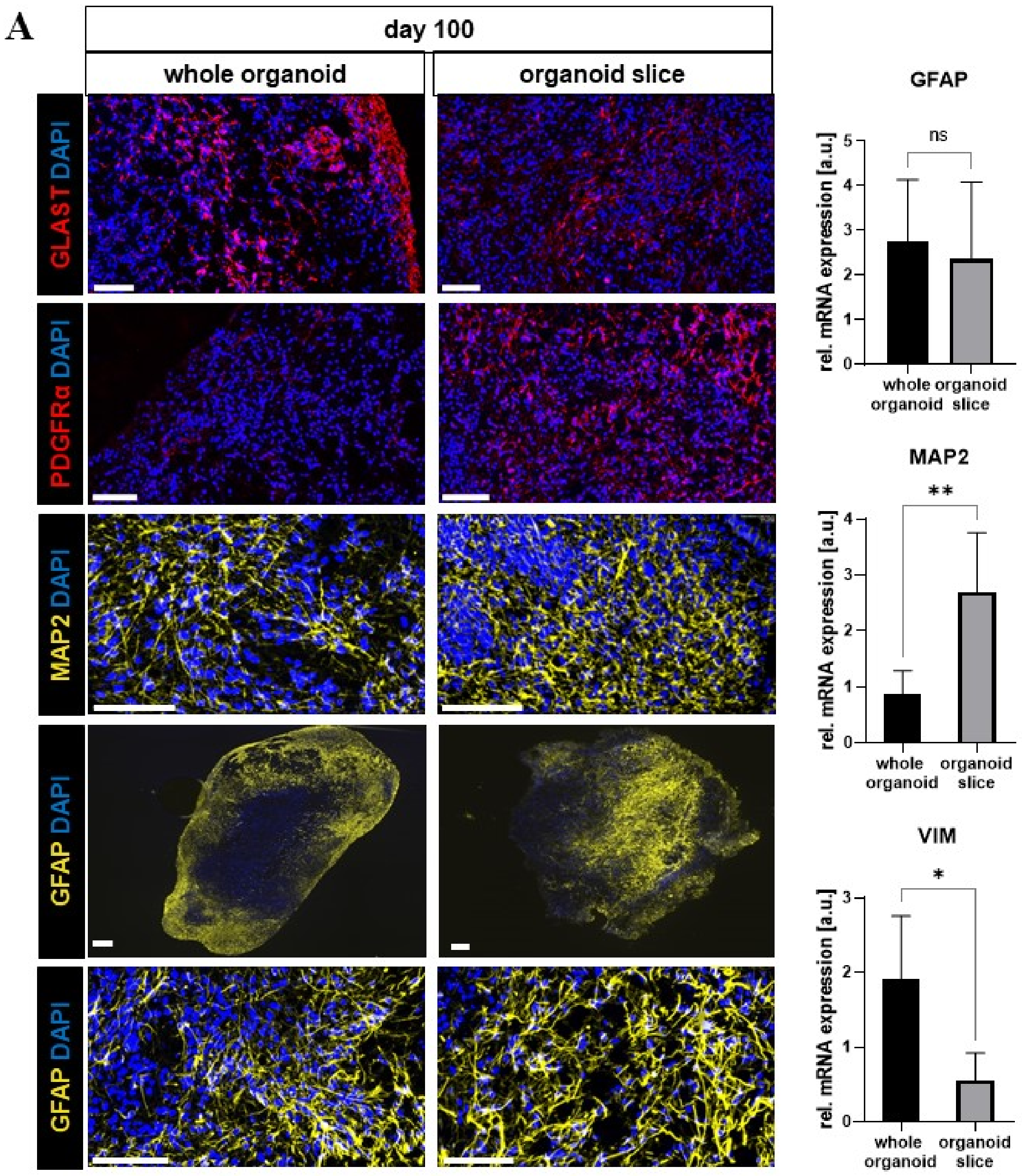
A
day 100
whole organoid
organoid slice
GLAST DAPI
PDGFRα DAPI
MAP2 DAPI
GFAP DAPI
GFAP DAPI
GFAP
ns
rel. mRNA expression [a.u.]
0
1
2
3
4
5
whole organoid
organoid slice
MAP2
**
rel. mRNA expression [a.u.]
0
1
2
3
4
whole organoid
organoid slice
VIM
*
rel. mRNA expression [a.u.]
0
1
2
3
whole organoid
organoid slice

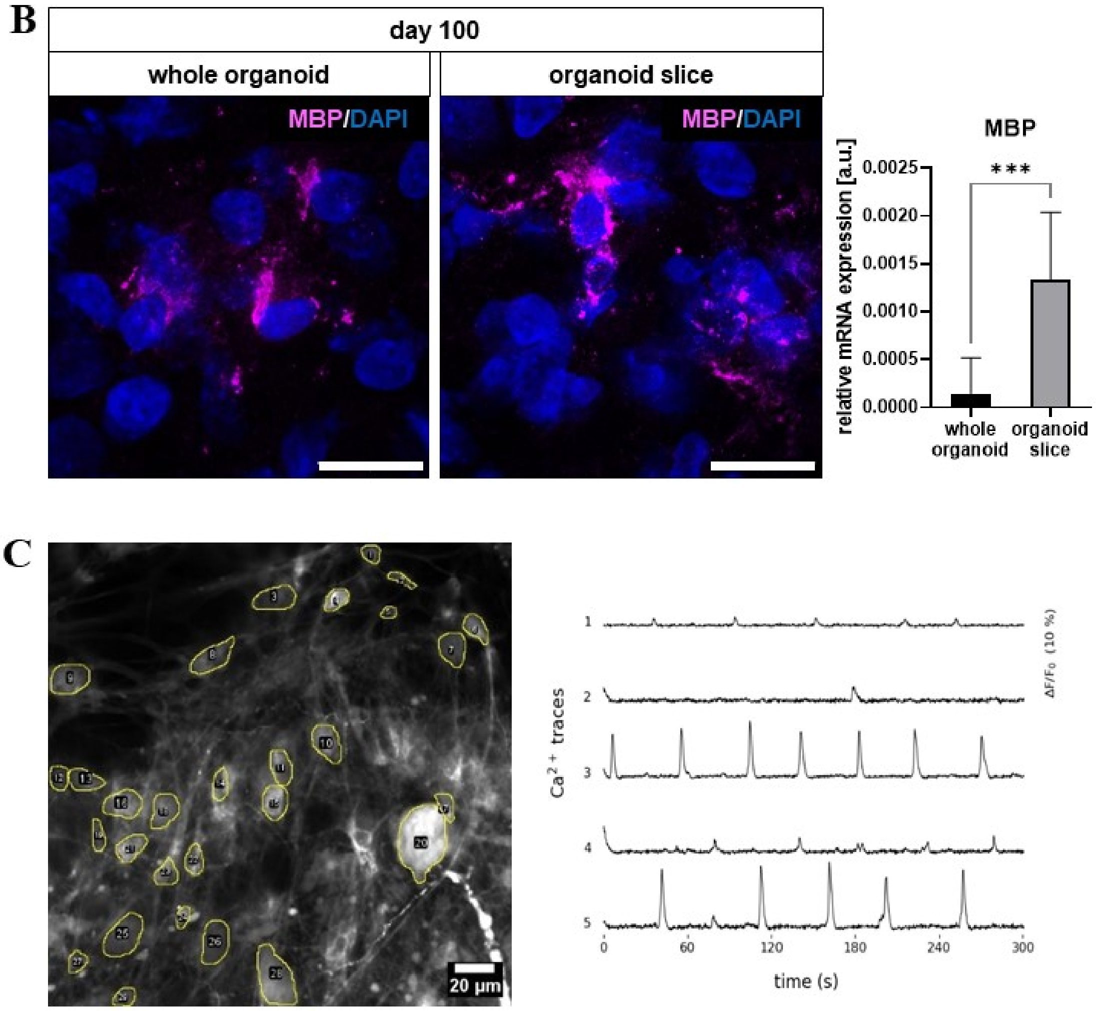


**Supplementary Figure 4. Maturation in whole organoids and organoid slices. A)** (left side) Representative immunofluorescence staining of GLAST (red), PDGFRα (red), MAP2 (yellow), and GFAP (yellow) for 100 days old whole organoids and organoid slices. Scale bar: 100 µm, 200 µm for GFAP in whole organoids. (right side) Relative mRNA expression of GFAP, MAP2, and VIM in 100 days old whole organoids and organoid slices. Data are presented as mean ± SD for five independent experiments (N = 5) and three organoids per experiment (n = 3) for whole organoids, and N = 6 (or N = 10 for VIM), n = 3 for organoid slices, * p<0.05, ** p<0.01, *** p<0.001, **** p<0.0001. Statistical analysis was done using Welch`s t-test. **B)** (top) Representative immunofluorescence staining of MBP (magenta) for 100 days old whole organoids and organoid slices. Scale bar: 20 µm. (bottom) Relative mRNA expression of MBP in 100 days old whole organoids and organoid slices. Data are presented as mean ± SD for seven independent experiments (N = 7) and three organoids per experiment (n = 3) for organoids, and N = 6, n = 3 for slices, * p<0.05, ** p<0.01, *** p<0.001, **** p<0.0001. Statistical analysis was done using Welch`s t-test. **C)** (left side) Representative image of Ca-imaging of multiple cell bodies (yellow circled) of one organoid slice. Scale bar: 20 µm. (right side) Exemplary Ca-traces of five different cells measured over 300 seconds.

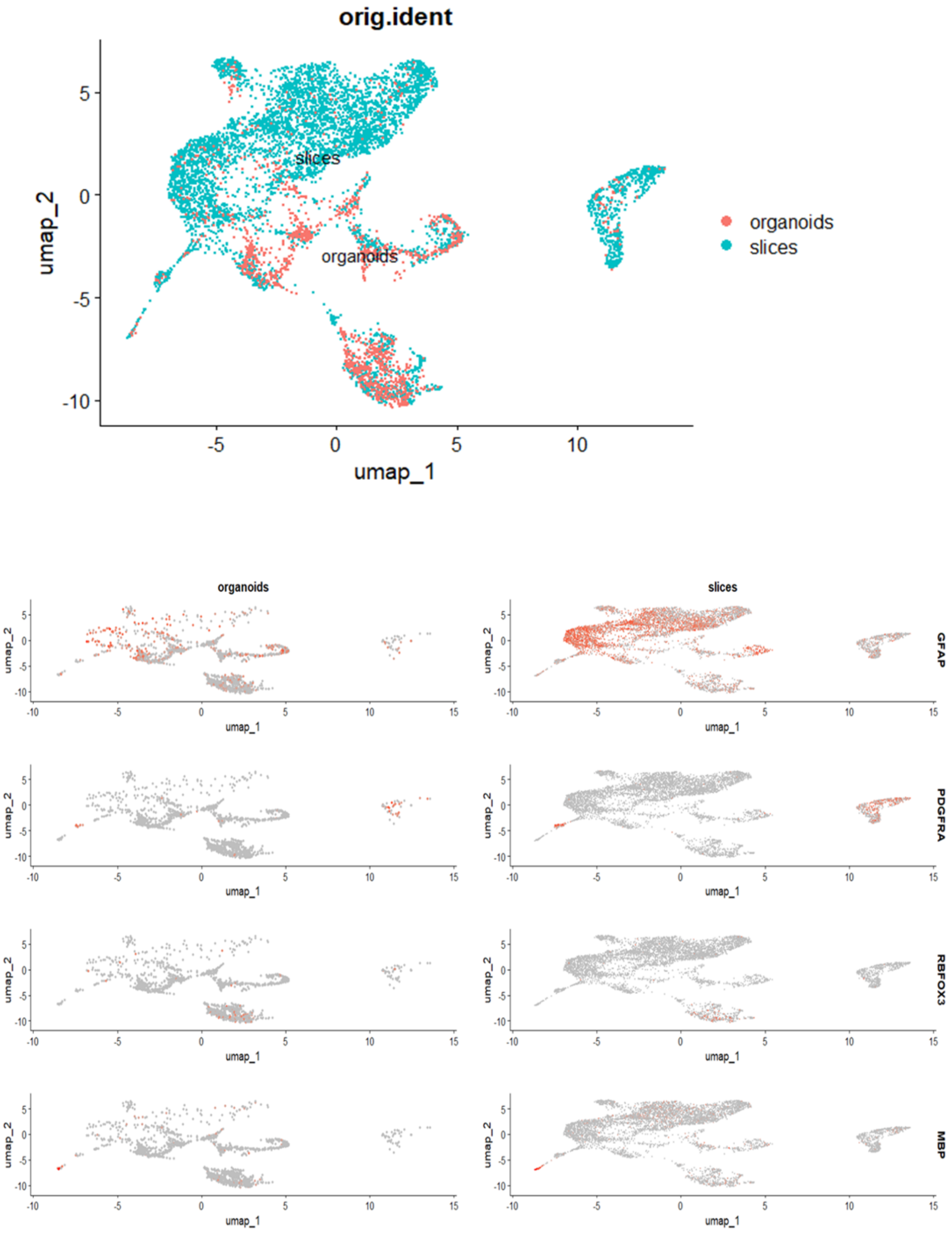


**Supplementary Figure 5. Whole organoids versus organoid slices at day 100 of the culture.** Single nuclei (sn) RNA analysis of integrated data sets for 100 days old whole organoids and organoid slices showing the distribution of GFAP, PDGFRα, RBFOX3, and MBP in single nuclei. One independent experiments (N = 1) with three organoids or slices (n = 3).

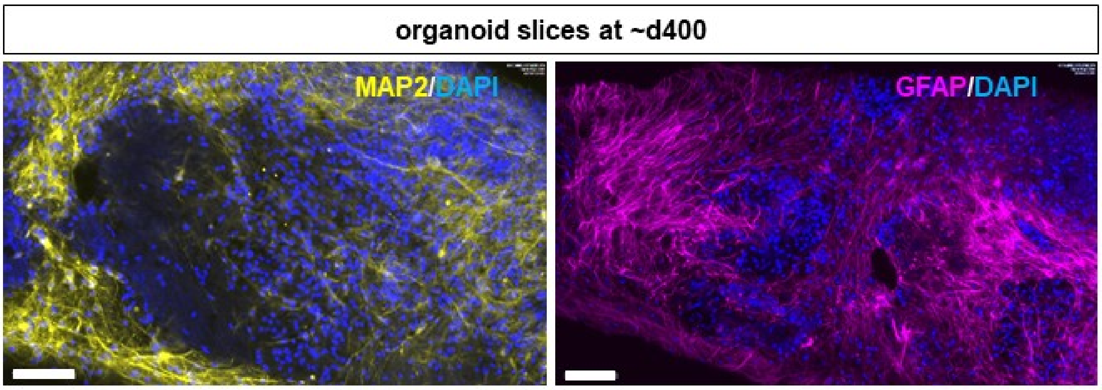


**Supplementary Figure 6**. **Organoid slices cultured over one year.** Representative immunofluorescence staining of MAP2 (yellow) and GFAP (magenta) for 400 days old organoid slices. Scale bar: 100 µm.

**A**

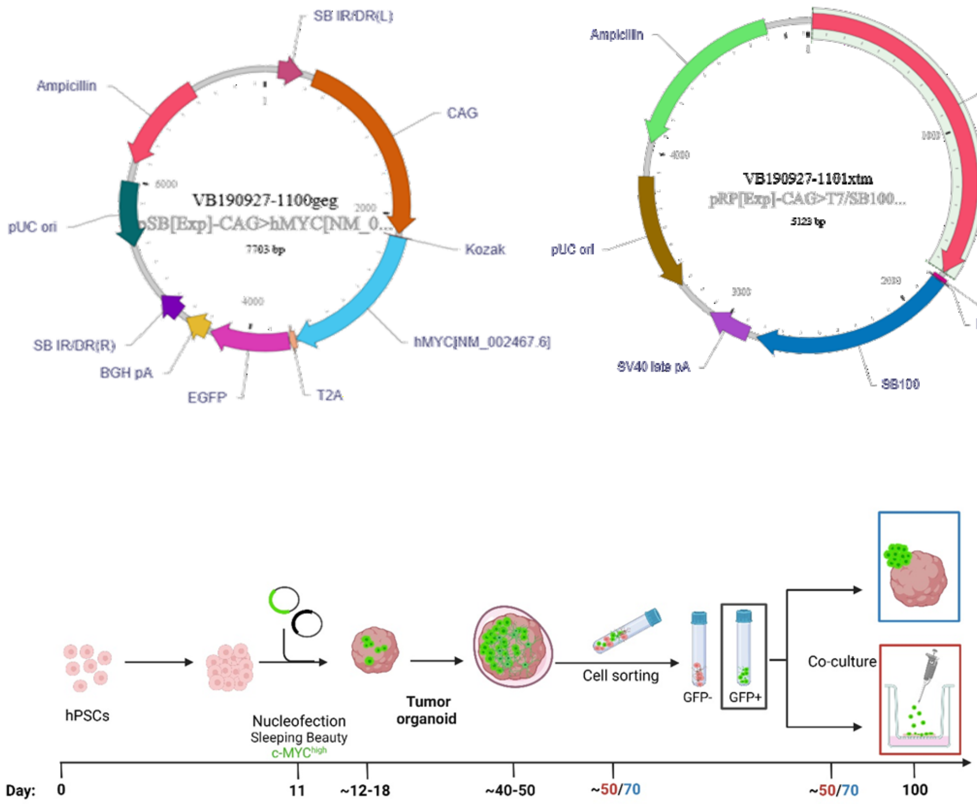

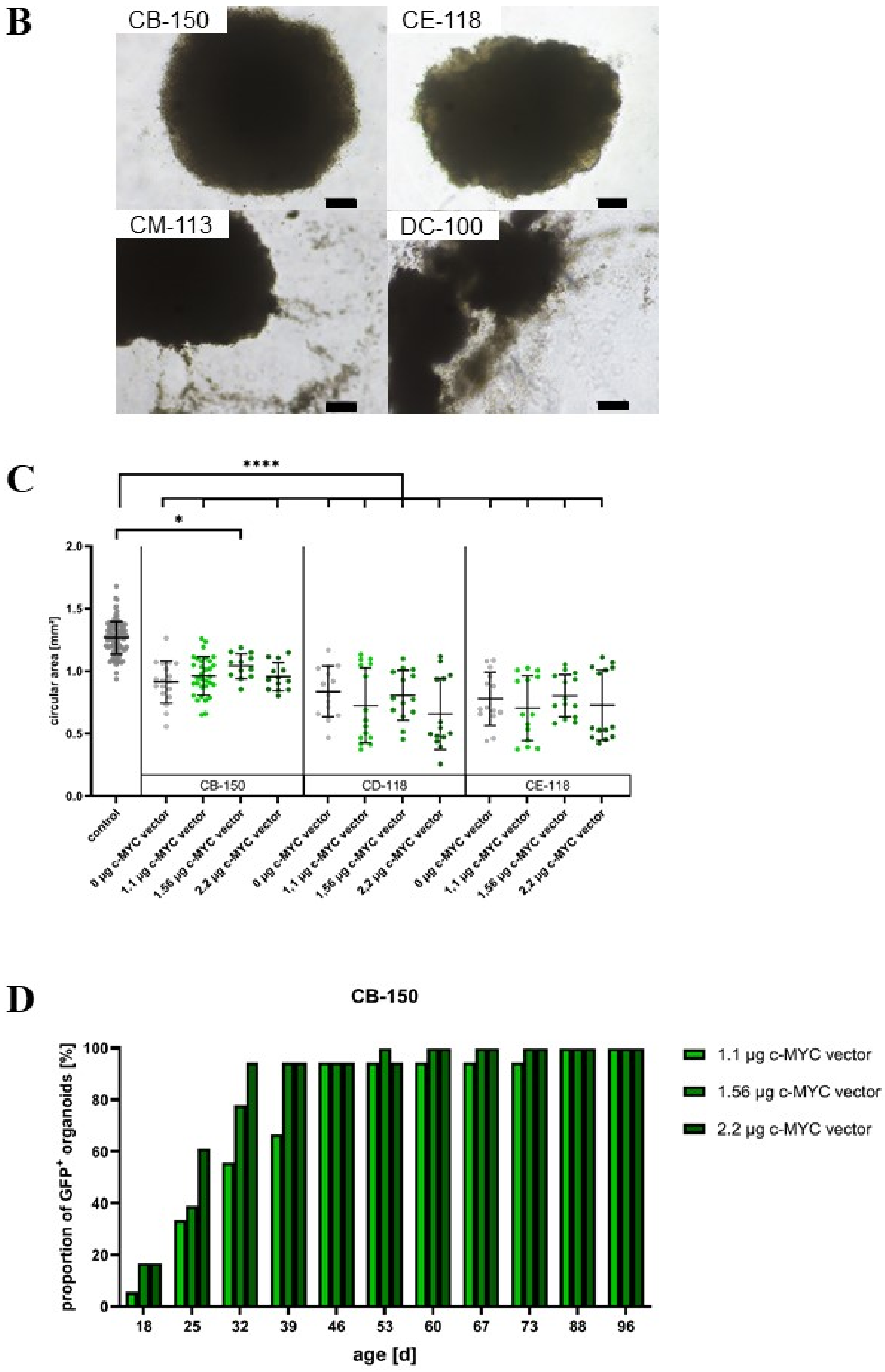
B
CB-150
CE-118
CM-113
DC-100
C
****
*
circular area [mm²]
2.0
1.5
1.0
0.5
0.0
CB-150
CD-118
CE-118
control
0 µg c-MYC vector
1.1 µg c-MYC vector
1.56 µg c-MYC vector
2.2 µg c-MYC vector
D
CB-150
proportion of GFP⁺ organoids [%]
100
80
60
40
20
0
18
25
32
39
46
53
60
67
73
88
96
age [d]
1.1 µg c-MYC vector
1.56 µg c-MYC vector
2.2 µg c-MYC vector

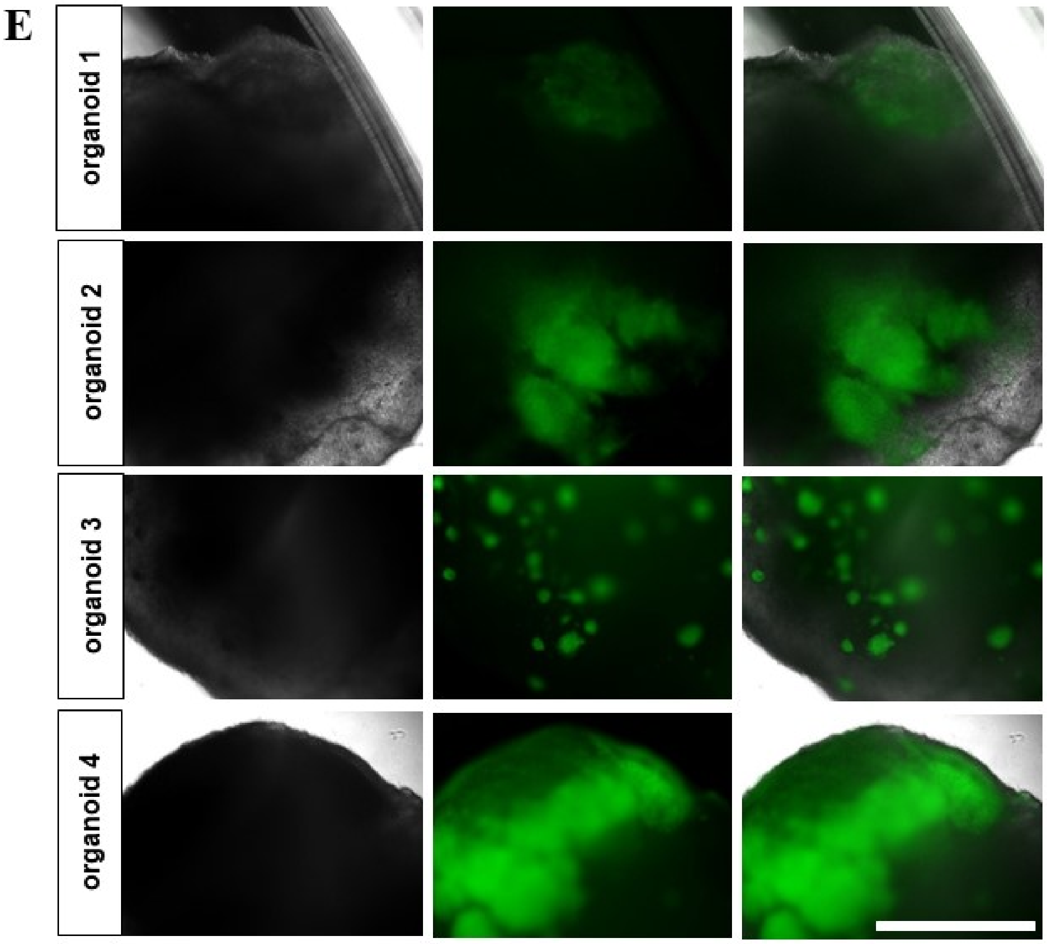


**Supplementary Figure 7. Establishment of the genetically modification protocol. A)** (top) Vector maps of and c-MYC (left, VB190927-1100geg) and Sleeping Beauty Transposase SB100 (right, VB190927-1101xtm). (bottom) Workflow of the genetically modification protocol for the generation of the tumor models. Embryoid bodies were generated using human pluripotent stem cells (hPSCs) and organoids were nucleofected on day 11 of culture using Sleeping Beauty Transposon System for overexpression of c-MYC oncogene with co-expressed GFP. $GFP^+$ genetically modified cells were detectable within one week and $GFP^+$ areas increased until isolation of genetically modified cells based on the presence of GFP signal using fluorescence-activated cell sorting at ~day 50 for co-culture with organoid slices or at ~day 70 for generation of tumor spheres to be assembled with whole organoids. Created with BioRender.com. **B)** Organoids 3 days after nucleofection using the program CB-150, CE-118, CM-113 or DC-100 of 4D-Nucleofector™ from Lonza. Scale bar 250 µm. **C)** Organoid size measured by the circular area 7 days after nucleofection (on day 18 of the culture) using the program CB-150, CD-188 or CE-118 of 4D-Nucleofector™ from Lonza. In each case, nucleofection was performed using 0 µg, 1.1 µg, 1.56 µg or 2.2 µg c-MYC vector and compared to non-nucleofected control organoids. Data are presented as mean ± SD for two to eight independent experiments (N = 2-8) and 2 to 30 organoids per experiment (n = 2-30), while each point represents the measurement of one single sample. * p<0.05, ** p<0.01, *** p<0.001, **** p<0.0001. Statistical analysis was done using Kruskal-Wallis Test with Dunn's post-test. **D)** Percentage of organoids showing a $GFP^+$ area after nucleofection with 1.1 µg, 1.56 µg or 2.2 µg c-MYC vector using the program CB-150 of 4D-Nucleofector™ from Lonza. Two independent experiments (N = 2) with nine organoids per experiment (n = 9). **E)** Exemplary $GFP^+$ areas of 4 different 53 days old organoids of one experiment. Scale bar: 1000 µm.

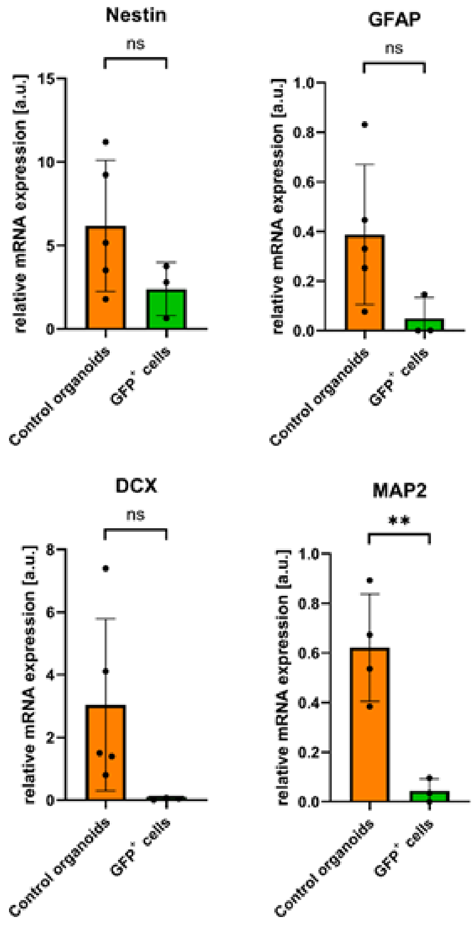


**Supplementary Figure 8. Genetically modified tumor-like cells generated in organoids show immature properties.** Relative mRNA expression of nestin, GFAP, MOG, DCX, and MAP2 in $GFP^{+}$ cells compared to whole non-nucleofected control organoids. Data are presented as mean ± SD for three independent experiments (N = 3) and one to two organoids per experiment (n = 1-2), * p<0.05, ** p<0.01, *** p<0.001, **** p<0.0001. Statistical analysis was done using unpaired t-test.

**Supplementary Table 1. Primary Antibodies**

| Antigen | Species | Company | Catalog No. | Dilution | RRID |
|---|---|---|---|---|---|
| SOX2 | rabbit | Thermo Fisher Scientific | A24339 | 1:100 | AB_2924437 |
| Nestin | mouse | BD Biosciences | 611658 | 1:500 | AB_399176 |
| DCX | rabbit | Abcam | ab18723 | | AB_732011 |
| CX43 | rabbit | Abcam | ab11370 | 1:1000 | AB_297976 |
| SMI312 | mouse | BioLegend | 837904 | 1:300 | AB_2566782 |
| GFP | chicken | Thermo Fisher Scientific | A10262 | 1:300 | AB_2534023 |
| VIM | mouse | Thermo Fisher Scientific | 14-9897-82 | 1:100 | AB_10597910 |
| CD133 | rabbit | Abcam | ab19898 | 1:200 | AB_470302 |
| c-Myc | mouse | Thermo Fisher Scientific | MA1-980 | 1:100 | AB_558470 |
| c-Myc | rabbit | Cell Signaling Technology | 13987 | 1:1000 | AB_2631168 |
| Ki-67 | rabbit | Abcam | ab16667 | 1:500 | AB_302459 |
| PDGFRapha | rabbit | Atlas Antibodies | HPA004947 | 1:250 | AB_2732399 |
| GLAST | rabbit | Thermo Fisher Scientific | PA5-111080 | 1:250 | AB_2856490 |
| MBP | rabbit | Abcam | ab218011 | 1:500 | AB_2895537 |
| SYN1 | rabbit | Synaptic Systems | 106103 | 1:500 | AB_11042000 |
| VAMP2 | rabbit | Synaptic Systems | 104202 | 1:1000 | AB_887810 |
| HOMER1 | mouse | Synaptic Systems | 160011 | 1:400 | AB_2120992 |

**Supplementary Table 2. Secondary Antibodies**

| Host | Target | Fluorophore | Company | Catalog No. | Dilution | RRID |
|---|---|---|---|---|---|---|

| | | | | | | |
|---|---|---|---|---|---|---|
| donkey | anti-mouse | AF488 | Thermo Fisher Scientific | A24350 | 1:250 | AB_2924437 |
| donkey | anti-rabbit | AF594 | Thermo Fisher Scientific | A24343 | 1:1000 | |
| goat | anti-mouse | AF568 | Thermo Fisher Scientific | A11004 | 1:1000 | AB_2534072 |
| goat | anti-rabbit | AF594 | Thermo Fisher Scientific | A11012 | 1:1000 | AB_2534079 |
| goat | anti-mouse | AF594 | Thermo Fisher Scientific | A21235 | 1:1000 | AB_2535804 |
| donkey | anti-mouse | AF594 | Thermo Fisher Scientific | A32744 | 1:1000 | AB_2762826 |
| goat | anti-chicken | AF647 | Thermo Fisher Scientific | A32933 | 1:1000 | AB_2762845 |
| goat | anti-mouse | AF647 | Thermo Fisher Scientific | A21235 | 1:1000 | AB_2535804 |

**Supplementary Table 3 Directly Labeled Antibodies**

| Antigen | Species | Company | Catalog No. | Dilution | RRID |
|---|---|---|---|---|---|
| MAP2 | rabbit | Abcam | ab225315 | 1:300 | AB_3517252 |
| GFAP | rabbit | Abcam | ab194325 | 1:300 | AB_3662092 |
| | | Abcam | ab216709 | 1:300 | AB_3662093 |
| Caspase-3 active | rabbit | GeneTex | GTX22302 | 1:300 | AB_384753 |

**Supplementary Table 4. Primers for qRT-PCR**

| Gene | Accession Nr. | Primer sequence (5´- 3´) |
|---|---|---|
| 18S rRNA | NR_003286.2 | ACTCAACACGGGAAACCTCACC (s) |
| | | CGCTCCACCAACTAAGAACGG (as) |

| | | |
|---|---|---|
| c-MYC | NM_002467.6/ NM_001354870.1 | TCGGATTCTCTGCTCTCCTC (s) |
| | | CCTGCCTCTTTTCCACAGAA (as) |
| p53 | NM_001407266.1 | CCTCAGCATCTTATCCGAGTGG (s) |
| | | TGGATGGTGGTACAGTCAGAGC (as) |
| Ki-67 | NM_002417.5 | GTGGTTCGACAAGTGGCCTT (s) |
| | | ACAACTCTTCCACTGGGACG (as) |
| NF1 | NM_000267.3 | GGACTCTAAGATCAACACCCTG (s) |
| | | CACCACACTCTGCACAATTCCAT (as) |
| PTEN | NM_001304718.2 | TGAGTTCCCTCAGCCGTTACCT (s) |
| | | GAGGTTTCCTCTGGTCCTGGTA (as) |
| CD133 | XM_054351160.1 | CACTACCAAGGACAAGGCGTTC (s) |
| | | CAACGCCTCTTTGGTCTCCTTG (as) |
| GLS | XM_054341407.1 | CAGAAGGCACAGACATGGTTGG (s) |
| | | GGCAGAAACCACCATTAGCCAG (as) |
| SNAI1 | NM_005985.4 | TGCCCTCAAGATGCACATCCGA (s) |
| | | GGGACAGGAGAAGGGCTTCTC (as) |
| MAP2 | NM_002374.3 | TGCGCTGATTCTTCAGCTTG (s) |
| | | TGTGTCGTGTTCTCAAAGGGT (as) |
| GFAP | NM_002055.5 | GTACCAGGACCTGCTCAAT (s) |
| | | CAACTATCCTGCTTCTGCTC (as) |
| VIM | NM_003380.5 | AGGCAAAGCAGGAGTCCACTGA (s) |
| | | ATCTGGCGTTCCAGGGACTCAT (as) |
| MBP | NM_001025081.1 | CTGTGCAACATGTACAAGGACTC (s) |
| | | GGGACAGTCCTCTCCCCTTT (as) |
| DCX | NM_178152.3 | AAGGACCTGTACCTGCCTCT (s) |
| | | TGAGCACTCTCCCCTCCTTT (as) |
